# DNA methylation signatures of frailty beyond age: a longitudinal study of female and male mice

**DOI:** 10.64898/2025.12.13.694109

**Authors:** Dantong Zhu, Shyleen Frost, Patrick Griffin, Maeve McNamara, David A. Sinclair, Alice E. Kane

## Abstract

Frailty is an age-related geriatric syndrome with largely unknown mechanisms. We conducted a longitudinal study of aging C57BL/6JNIA mice (females; *n* = 40, male; *n* = 49), measured frailty index and derived DNA methylation data from PBMCs. We selected frailty-related differentially methylated CpGs and determined differentially methylated regions (DMRs), focusing on both age-independent and -dependent frailty, and using both mixed-sex and sex-stratified subgroups. We propose a joint set of 925 frailty-related DMRs, perform an association study with frailty outcomes, build epigenetic frailty clocks and validate in mice with interventions. Notably, age-independent frailty DMRs are enriched in nervous and endocrine pathways, distinct from signaling and lipid metabolism pathways identified from age-dependent DMRs. We observe hypermethylation in signaling pathways and hypomethylation in lipid metabolism and cytochrome P450 pathways with frailty progression. 36 DMRs show consistent associations in validation. These findings highlight distinct epigenetic signatures underlying frailty and aging, with potential sex-specific mechanisms.

## Introduction

Aging is highly heterogeneous, with substantial variability in health and function amongst individuals of the same age^1^. Frailty captures such variability and reflects the overall decline in health with age^2^. Frailty index (FI) is utilized to measure frailty and represents the proportion of age accumulated health-related deficits present in an individual^3,4^. Higher FI values indicate a greater degree of frailty and are associated with an increased susceptibility to diseases and mortality^4,5^. Frailty indices have been adopted for use in other mammals, including mice^6^. Currently, there are no accepted frailty biomarkers^7^, and the underlying molecular mechanisms of frailty, especially those distinct from aging itself, remain largely unknown. The identification of frailty biomarkers and determinants would allow for earlier recognition and tracking of frailty over time, testing of frailty interventions and possible discovery of druggable frailty targets^7^.

DNA methylation (DNAm) represents an attractive biomarker option as it changes predictably with age and disease, can be measured across sample types, can provide insight into biological pathways^8^ and appears to be responsive to interventions^9^. DNA cytosine methylation, which is predominantly maintained at CpG loci, plays a central role in regulating gene expression and chromatin structure. Studies of the associations of DNAm at these specific CpG sites with aging cover a wide range of developmental stages, tissues, and animal species. Notably, epigenetic clocks, algorithms that predict chronological age or other health outcomes from methylation patterns at 10-100s of individual CpGs have been developed, including Horvath’s clock^10^, GrimAge^11^ and DunedinPACE^12^. Differentially methylated regions (DMRs) are fragments of the genome that harbor multiple methylation loci^13^, and are regarded as functional regions involved in gene transcription regulation.

Studies in humans have unravelled age-related methylation changes linked to inflammaging^14^ and immune systems^15,16^, as well as metabolic disorders^17,18^. Age-related methylation changes are also observed in other mammalian animals e.g. mice and monkeys^19,20^. While most studies are cross-sectional, longitudinal analyses have revealed dynamic epigenetic changes over time, including variably methylated positions (VMPs) that reflect inter-individual variability linked to aging and environmental factors^21^. Epigenetic entropy measures, such as Shannon entropy, further quantify methylation complexity, offering additional insight into aging-related epigenetic drift^22^. These findings are consistent with the Information Theory of Aging, which proposes that aging results from the progressive loss of epigenetic information, a process that may be reversible^23^. Together, these approaches provide complementary views of the temporal and stochastic aspects of epigenetic aging, although their application to frailty remains limited. Associations of frailty with global DNAm changes have been investigated, with emerging evidence suggesting a decrease in global DNAm with frailty progression^24^. However, specific frailty-related CpGs and DMRs have not been extensively explored^25,26^, and inter-individual variability in DNA methylation with frailty remains largely unexamined.

Additionally, most prior studies of DNA methylation in aging or frailty focus on one sex or include sex as a covariate, with limited investigation into sex differences in frailty- and age-related methylation. Yet, females and males present distinct epigenetic profiles^27,28^, and the differences appear to persist throughout aging^29,30^. Sex-specific methylation patterns with aging have been reported at individual loci—for example, lower methylation levels of *PRR4* regulating loci^30,31^ are observed in males. Although a few studies using epigenetic clocks have also hinted at sex differences in frailty^32,33^, nevertheless they do not dive into specific loci nor the underlying mechanisms.

In this study, we conducted a longitudinal study of female and male mice and generated matched DNAm profiles and frailty measurements across five time points. We used differential methylation analysis to select differentially methylated positions (DMPs) and then identified differentially methylated regions (DMRs). At the DMR level, we selected candidate methylation features with a focus on age-independent and -dependent components of frailty, in both mixed-sex and -specific subgroups. We generated a joint set of frailty-related DMRs, whereupon we performed an association study, built an epigenetic frailty clock and validated in mice with interventions. Our findings show that the age-independent frailty-related DMRs are enriched in insulin resistance, different from the lipid metabolism and signaling pathways identified from the age-dependent frailty-related DMRs. We observe sex dimorphisms in frailty-related DMRs, with distinct pathway enrichments in females and males. Overall, our study provides candidate methylation features linked to frailty beyond age, offering potential biomarkers for clinical validation and shedding light on possible mechanisms underlying sex differences in frailty and aging.

## Results

### DNA methylation array data variation

We performed a longitudinal study of female (*n* = 40) and male (*n* = 49) C57BL/6JNIA mice at up to 5 time points, and derived a total of 315 samples that have DNAm array data and FI measurement (**Supplementary Table 1**; **Supplementary Fig.1**). In order to investigate DNAm and mechanisms in naturally aging mice, we used untreated mice (female, *n* = 20; male, *n* = 25), including 153 samples, as the discovery cohort (**Fig.1**). We obtained 242,599 CpG sites (CpGs) (85.0% of 285,255 CpGs included in the array) after coverage and variation filtering (see Methods). Global DNAm levels (beta values) decreased with aging in both males and females (**Supplementary Fig.2a**). As expected, the majority of DNAm beta values are concentrated at the extremes, with values clustering near 0 or 1. While CpGs with low methylation levels (close to zero) showed a similar distribution, those with high methylation levels (close to 1) displayed varying density levels across age (**Supplementary Fig.2b**). The results suggest that epigenetic aging is primarily driven by alterations in highly methylated regions.

**Fig.1.**
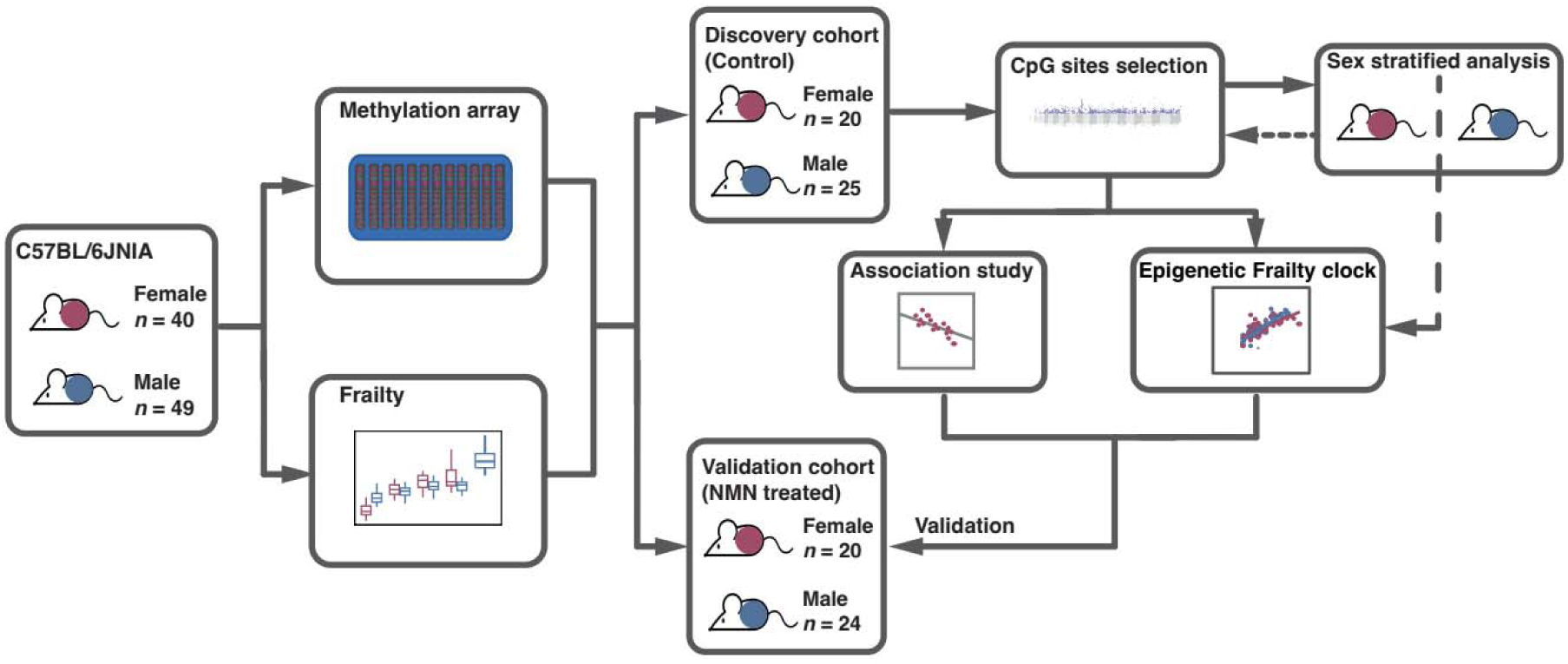
Schematic diagram of the analysis workflow.

To further investigate the variations in CpG methylation levels across samples, we performed a principal component analysis (PCA) using M-values (logit transformation of beta values that are more statistically valid) of the 242,599 CpGs. The PCA plots indicate separation of samples across ages and by sex (**Supplementary Fig.2c**), as well as by log-transformed FI (logFI, to reduce skewness and improve model fit) (**Supplementary Fig.2d**). Variance partitioning analysis on the top 121 PCs that explained more than 95% of data variance revealed that sex explained approximately 10.5% of the overall methylation variance, and mouse ID (i.e. variance within a single mouse over time), age and FI residuals (age factor excluded) explain 9.8%, 5.0% and 1.1% of the variance, respectively (**Supplementary Fig.2e**). Altogether, the results indicate the dynamics of DNA methylation are strongly associated with individual variation and sex, but also associated with age and frailty.

### DNA methylation signatures of frailty

To identify DNAm loci that are related to frailty, we performed an Epigenome-Wide Association Study (EWAS) with logFI as the outcome, including sex as a fixed effect and mouse ID as a random effect to account for repeated measures. We found 61,112 differentially methylated positions (DMPs) associated with FI, representing 25.2% of the 242,599 CpGs included in the analysis (**Fig.2a**). These DMPs are distributed across the genome, including the sex chromosomes, with 2,184 DMPs in X chromosome (accounting for 21.1% of CpGs on X included in the EWAS) and 10 DMPs in Y chromosome (10.6% of CpGs on Y) (**Supplementary Fig.3a**). When examining the genomic locations of these DMPs, a smaller proportion of DMPs are located within CpG islands (χ²(1) = 730.29, *p* < 0.001) and TSS200 regions (χ²(1) = 547.83, *p* < 0.001) than across all CpGs considered (**Supplementary Fig.3b**), suggesting that frailty-associated methylation changes are more likely to occur outside traditional CpG islands and core promoters. Given that DNA methylation changes are frequently regional rather than restricted to individual CpG sites, we identified 1,882 differentially methylated regions (DMRs) that capture coordinated epigenetic changes by aggregating information from neighboring CpGs (see Methods) (**Supplementary Fig.3c**). These DMRs contain a total of 12,543 CpGs, including 5,691 DMPs and 6,852 non-DMPs. We restricted subsequent analyses to those CpGs within DMRs because regional methylation changes are more biologically meaningful and less prone to noise than isolated CpG signals^34^.

**Fig.2.**
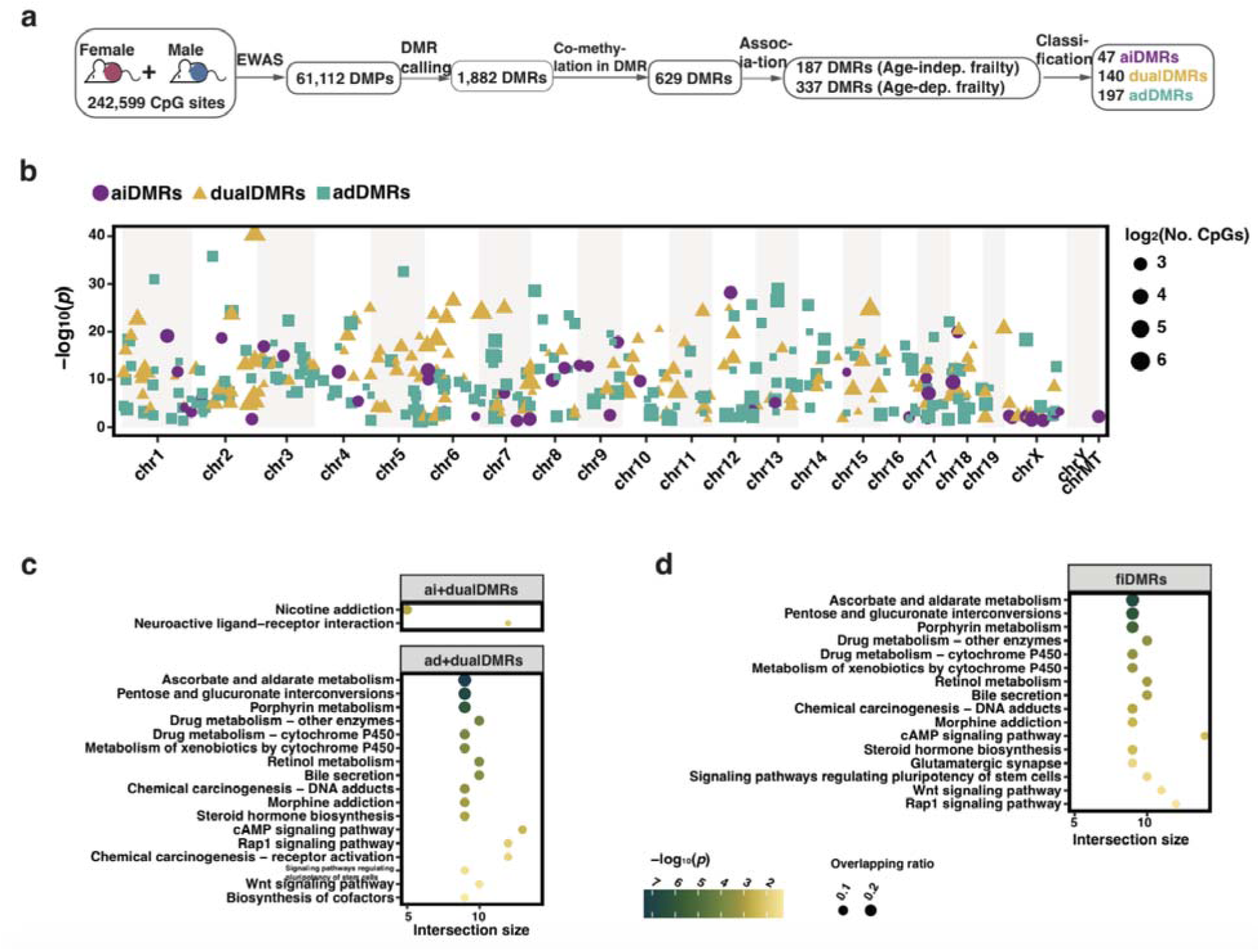
Selection, classification, and functional analysis of frailty-related methylation markers in the sex-inclusive subgroup. **a.** Schematics diagram of the DMR selection process. From differentially methylated sites (DMPs), differentially methylated regions (DMRs) were determined, and then classified based on associations with age-independent (Age-indep.) and age-dependent (Age-dep.) frailty as aiDMRs (only associated with Age-indep. frailty), adDMRs (only associated with Age-dep. frailty) and dualDMRs (simultaneously associated with above both frailty). **b.** Manhattan plot displaying frailty index-related DMRs (fiDMRs). Each DMR is colored and shaped according to its classification. Point size reflects the number of CpGs. **c.** Probe set enrichment analysis of pathways for CpGs within corresponding DMRs, aiCpGs + dualCpGs for age-indep. frailty and adCpGs + dualCpGs are for age-dep. frailty. **d.** Probe set enrichment analysis of pathways for fiDMRs.

To distinguish methylation features specific to frailty from those driven by chronological aging, we determined epigenetic changes associated with the age-independent and age-dependent components of frailty (hereafter referred to as AIFI and ADFI) at the DMR level (**Fig.2a**). After excluding DMRs with low co-methylation levels^35^ to reduce noise from random methylation variation, we obtained 629 DMRs (no. of CpGs ≥ 5). We tested the associations of these 629 DMRs with frailty using linear mixed models, and found 187 and 337 DMRs were associated with AIFI and ADFI, respectively (**Supplementary Fig.3d**). According to the association with frailty, we then determined 47 aiDMRs (DMRs that were only AIFI-related), 140 dualDMRs (DMRs that were simultaneously related to AIFI and ADFI) and 197 adDMRs (DMRs that were only ADFI-related) (a total of 384 frailty-related DMRs; fiDMRs) that contained 335, 1,514 and 1,488 CpGs respectively (a total of 3,337 frailty-related CpGs; fiCpGs) (**Fig.2b**; **Supplementary Table 2**).

Probe set enrichment analysis was performed for these DMRs to identify overrepresented genes and pathways. DMRs associated with AIFI (aiDMRs + dualDMRs) were linked to 412 genes, including *Pou3f3*, *Pantr1*, *Foxg1*, *Pantr2*, *Sfmbt2* and *Nr2f2* (**Supplementary Table 3**). Representative examples for DMRs overlapping the *Foxg1* and *Pou3f3* genes are shown in **Supplementary Fig.3e** and **f**, where the increasing methylation levels across these regions with increasing frailty scores can be clearly visualised. DMRs associated with AIFI were enriched in neuroactive ligand-receptor interaction and nicotine addiction pathways (**Fig.2c**). CpGs within DMRs associated with ADFI (adDMRs + dualDMRs) presented dynamic patterns in aging (**Supplementary Fig.3g**) and were found predominantly enriched in lipid metabolism and signaling related pathways (**Fig.2c**). When considering all 384 fiDMRs, pathway analysis indicated 16 pathways enriched, including retinol metabolism, morphine addiction, glutamatergic synapse, and Wnt signaling pathway (**Fig.2d**). Taken together, our findings identified fiDMRs from AIFI and ADFI are associated with lipid metabolism, insulin resistance, nervous system, and signaling pathways.

### Sex differences in frailty-related DNA methylation signatures

Sex dimorphisms in frailty and aging are widely observed, hence sex specific CpGs that are related to frailty are of interest. We applied a sex stratification analysis approach to identify candidate frailty-related methylation features for males and females. We performed EWAS for FI within each sex to identify sex-specific DMPs, and detected 90,369 and 69,091 DMPs associated with frailty in females and males, respectively (**Supplementary Fig.4a**). For both males and females, proportions of DMPs in CpG islands and close to TSSs were lower than expected, when compared to the total 242,599 CpGs included in our analysis. Interestingly, females had higher proportions of DMPs in these locations than males (**Supplementary Fig.4b**).

When considering the sex chromosomes specifically, there were 4,464 (43.2% of those measured on X) DMPs detected in the X chromosome in females. For males, 2,702 (26.1%) DMPs were X chromosome-linked and 4 (4.3%) DMPs were Y chromosome-linked. We also observed sex differences in the distribution of methylation levels at the X chromosome across all CpGs subjected to analysis including all ages (**Supplementary Fig.4c**). Additionally, we identified CpGs that were likely undergoing X chromosome inactivation (XCI) and inactivation escape (i.e. activation, XCA) in females by applying a validated decision tree, comparing methylation levels across all measured CpGs on the male and female X chromosomes^36^. We identified 1,018 (9.8% of 10,338 CpGs measured on X chromosome) XCI related CpGs and 100 (1.0%) XCA related CpGs (**Supplementary Fig.4d**). Notably, 13 of these XCA related CpGs were also identified as frailty-related DMPs, suggesting that X chromosome inactivation escape may contribute to frailty for females. Probe enrichment analysis indicates these 13 CpGs were linked to *Slitrk4,* a transmembrane protein with roles in neuronal functioning, and *Bclaf3*, a DNA binding protein.

We then dissected sex specific methylation features associated with AIFI and ADFI, using the same analysis pipeline as for the sex-inclusive subgroup above, stratifying by sex. From the 90,369 DMPs identified in females, we identified 3,280 DMRs harboring 21,488 CpG sites (including 9,767 DMPs) (**Fig.3a; Supplementary Fig.5a**). We selected 21 dualDMRs and 750 adDMRs (a total of 771 fiDMRs) that contain 468 dualCpGs and 5,623 CpGs (a total of 6,091 fiCpGs), but no sites were solely associated with age-independent frailty index (**Fig.3b**; **Supplementary Fig.5b**; **Supplementary Table 4**). When considering dualDMRs (those associated in both age independent and dependent frailty analyses), we detected 66 genes including *GNAS* (**Supplementary Fig.5c**), *GNAS-AS1* and *Igf2r* (**Supplementary Table 5**). DMRs presenting ADFI signals, also female fiDMRs in females, were enriched for predominantly signaling pathways and neuronal activity (**Fig.3c**).

**Fig.3.**
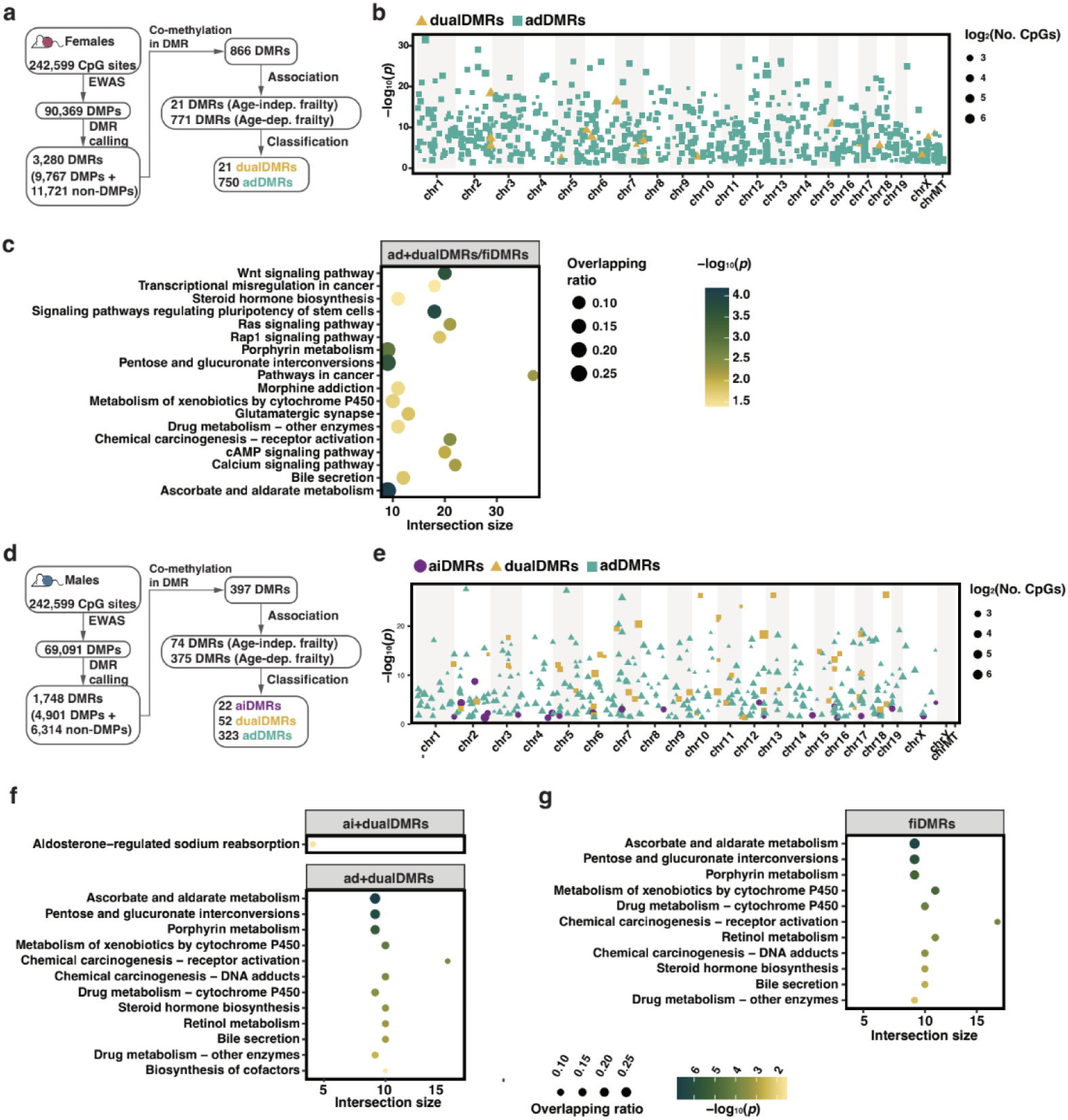
Selection, classification, and functional analysis of frailty-related methylation markers in females and males. **a.** Schematics diagram of the DMR selection process in females. From differentially methylated sites (DMPs), differentially methylated regions (DMRs) were determined, and then classified based on associations with age-independent (Age-indep.) and age-dependent (Age-dep.) frailty as aiDMRs (only associated with Age-indep. frailty), adDMRs (only associated with Age-dep. frailty) and dualDMRs (simultaneously associated with above both frailty). **b.** Manhattan plot displaying fiDMRs in females. Each DMR is colored and shaped according to its classification. Point size reflects the number of CpGs. **c.** Probe set enrichment analysis of pathways for CpGs within fiDMRs in females. **d.** Schematics diagram of the DMR selection process in males. **e.** Manhattan plot displaying fiDMRs in males. Each DMR is colored and shaped according to its classification. Point size reflects the number of CpGs. **f.** Probe set enrichment analysis of pathways for CpGs within corresponding DMRs in males. ‘aiCpGs + dualCpGs’ represents age-indep. frailty, adCpGs + dualCpGs for age-dep. frailty. **g.** Probe set enrichment analysis of pathways for CpGs within fiDMRs in males.

For males, from the 69,091 DMPs, we identified 1,748 DMRs that contain 11,215 CpG sites (**Fig.3d; Supplementary Fig.5d**). 22 aiDMRs, 52 dualDMRs and 323 adDMRs (a total of 3,000 fiCpGs) were selected (**Fig.3e**; **Supplementary Fig.5e**; **Supplementary Table 6**). We found AIFI DMRs are associated with 189 genes (**Supplementary Table 7**), for instance *Il27* and *Pik3ip1*, and enriched in aldosterone-regulated sodium reabsorption (**Fig.3f**). ADFI DMRs were found predominantly enriched in lipid metabolism and cytochrome P450 pathways (**Fig.3f**). Regarding all 397 male fiDMRs, 11 pathways were found, including bile secretion and metabolism of xenobiotics by cytochrome P450 (**Fig.3g**).

### Comparative analysis of sex-specific and common fiDMRs

From the sex stratified analysis, we obtained 771 female- and 397 male-fiDMRs. Of the total 903 DMRs, 265 DMRs (harboring 2,075 fiCpGs) were common between females and males (**Fig.4a**; **Supplementary Table 8**). Notably, while all 265 DMRs were classified as age dependent (adDMRs) in females, 9 and 27 were classified as aiDMRs and dualDMRs in males. These 36 (9+27) DMRs are enriched in Fc gamma R-mediated phagocytosis, suggesting possible sex differences in this pathway in frailty. The remaining 229 DMRs that were simultaneously associated with ADFI in both sexes were enriched in predominantly signaling and lipid metabolism pathways (**Fig.4b**), as seen in our mixed sex analysis. 506 female-specific DMRs (example shown in **Supplementary Fig.5f**), were either positively (*n* = 309) and negatively (*n* = 197) associated with frailty, and were enriched in mainly signaling pathways and neuronal function, respectively (**Fig.4c**). Of the 132 male-specific DMRs, 46 were positively associated and 86 negatively associated with frailty (example in **Supplementary Fig.5g**), and were linked to 99 (e.g. *Akt1s1*) and 231 (e.g. *Il27*) genes respectively.

**Fig.4.**
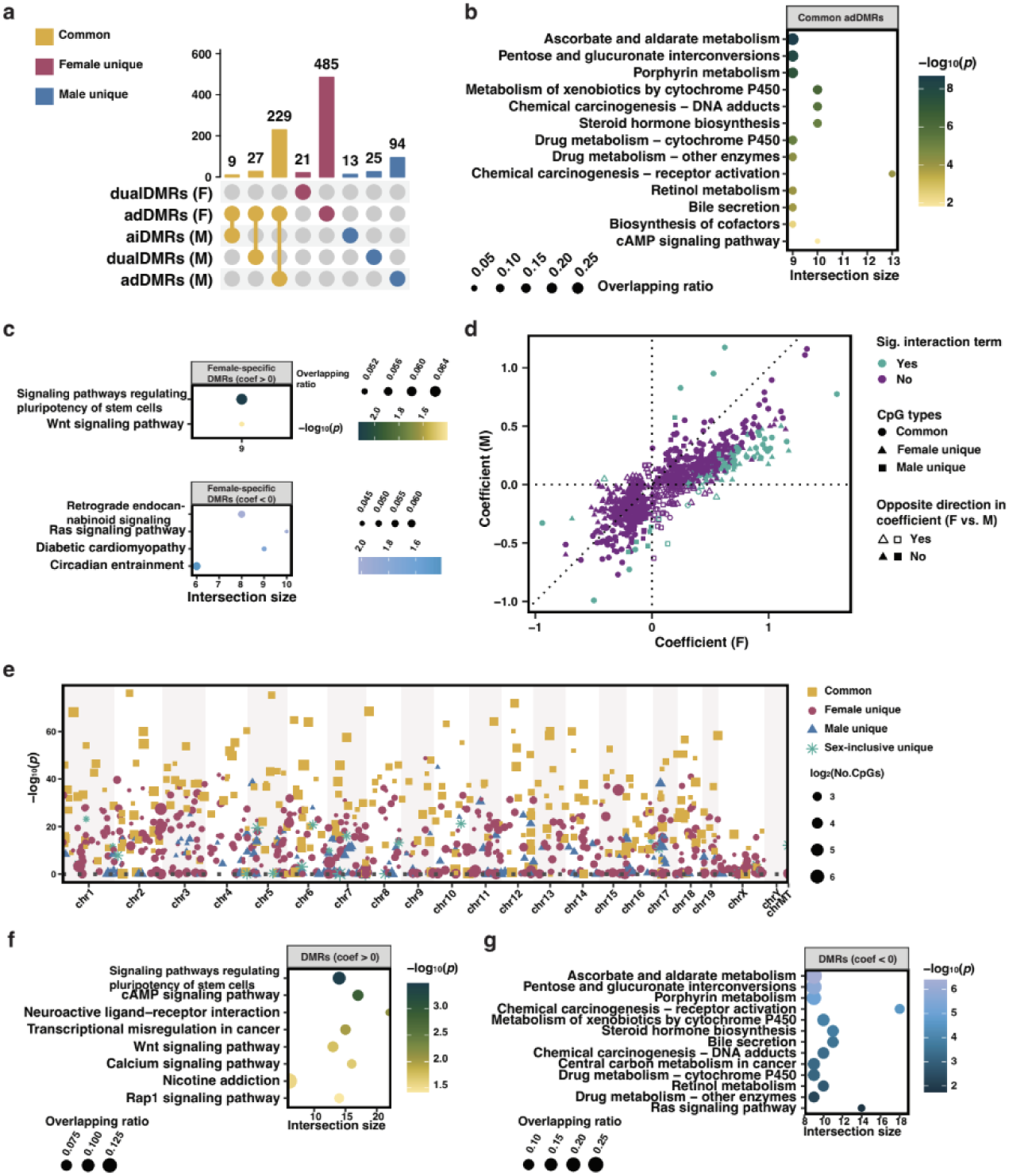
Sex-specific and shared frailty-related methylation signatures. **a.** Upset plot showing sex difference in frailty-related differentially methylated regions (fiDMRs). Within each sex, fiDMRs were classified according to the association with frailty traits, with aiDMRs for DMRs that are only associated with age-independent frailty, adDMRs for those only associated with age-dependent frailty and dualDMRs for those associated with both traits. **b.** Probe set enrichment of common adDMRs (shared by females and males). **c.** Probe set enrichment analysis of DMRs found in females stratified by positive and negative effect size of associations with frailty. **d.** Scatter plot showing sex-specific effect size of fiDMRs. Each point represents one DMR derived from the union of female- and male-specific fiDMRs. For each DMR, effect sizes were estimated separately in females and males using linear mixed models, and classified based on whether the directions of effect were the same or opposite between sexes. The significance of the frailty by sex interaction term was determined in the whole cohort (females and males combined) analysis. **e.** Manhattan plot showing combined fiDMRs derived from analyses within sex-inclusive and -specific subgroups. Each DMR is colored and shaped based on its classification according to subgroups in which the DMR was found. Point size reflects the number of CpGs. **f.** Probe set enrichment analysis of pathways for CpGs within DMRs that exhibit positive effect size in frailty progression. **g.** Probe set enrichment analysis of CpGs within DMRs that exhibit negative effect size in frailty progression.

To further investigate sex-specific differences in methylation changes, we performed sex interaction analysis in the combined female and male samples and derived effect sizes of these DMRs in females and males, respectively. Interestingly, we found 460 (90.1% of 506) female-fiDMRs and 120 (90.1% of 132) male-fiDMRs had non-significant sex interaction terms, indicating that in this analysis these CpGs were not statistically different between the sexes in terms of their association with frailty, despite being identified as sex specific in our stratified analysis. Overall, **Fig.4d** shows that, despite being identified in only one sex, the majority (796, 88.2%) of DMRs show the same direction of association of methylation with frailty across both sexes. Of particular interest, among the 107 DMRs showing the opposite direction in effect size with frailty in females and males, 6 were simultaneously identified in females and males as fiDMRs, within which the CpGs were linked to 18 genes, including *Angptl2*, suggesting these genes undergo sex-biased regulation of expression in frailty progress.

Although sex differences in epigenetic signatures are clear, there is also substantial overlap in the biological mechanisms underlying frailty across sexes. Therefore, integrating DMRs identified from both sex-inclusive and sex-stratified analyses provides a more comprehensive representation of the DNAm frailty signature. We identified 925 candidate fiDMRs that are associated with frailty in 1) both males and females, 2) females only, 3) males only or 4) only when males and females are combined for sex-inclusive analysis (**Fig.4e**; **Supplementary Table7**). Across fiDMRs, 526 (including 4,155 fiCpGs) were positively associated with frailty, where they gained methylation with frailty progression, and were linked to 1,041 genes (e.g. *Prkcz*) and predominantly enriched in signaling pathways (**Fig.4f; Supplementary Fig.6a**). 399 DMRs were negatively associated with frailty i.e. hypomethylation in frailty progression, and were enriched mainly in lipid metabolism and cytochrome P450 related pathways (**Fig.4g**; **Supplementary Fig.6b**).

Taken together, our findings confirm sex-inclusive analysis with the identification of common fiDMRs across sexes enriched in signaling and lipid metabolism pathways, whilst also revealing some sex differences in methylation patterns.

### DMRs associated with current and future frailty

While we have investigated the associations between methylation changes and frailty at the same time point, we were interested in assessing the relationship between identified DMRs and subsequent frailty outcomes aiming to evaluate the utility of these DMRs as prospective biomarkers of frailty risk. We focused on future FI outcomes and applied linear mixed models using each of these outcomes as presented in **Fig.5a**. Results are summarized in **Supplementary Table 9**, and DMRs showing a significant interaction with age in corresponding analyses were excluded.

**Fig.5.**
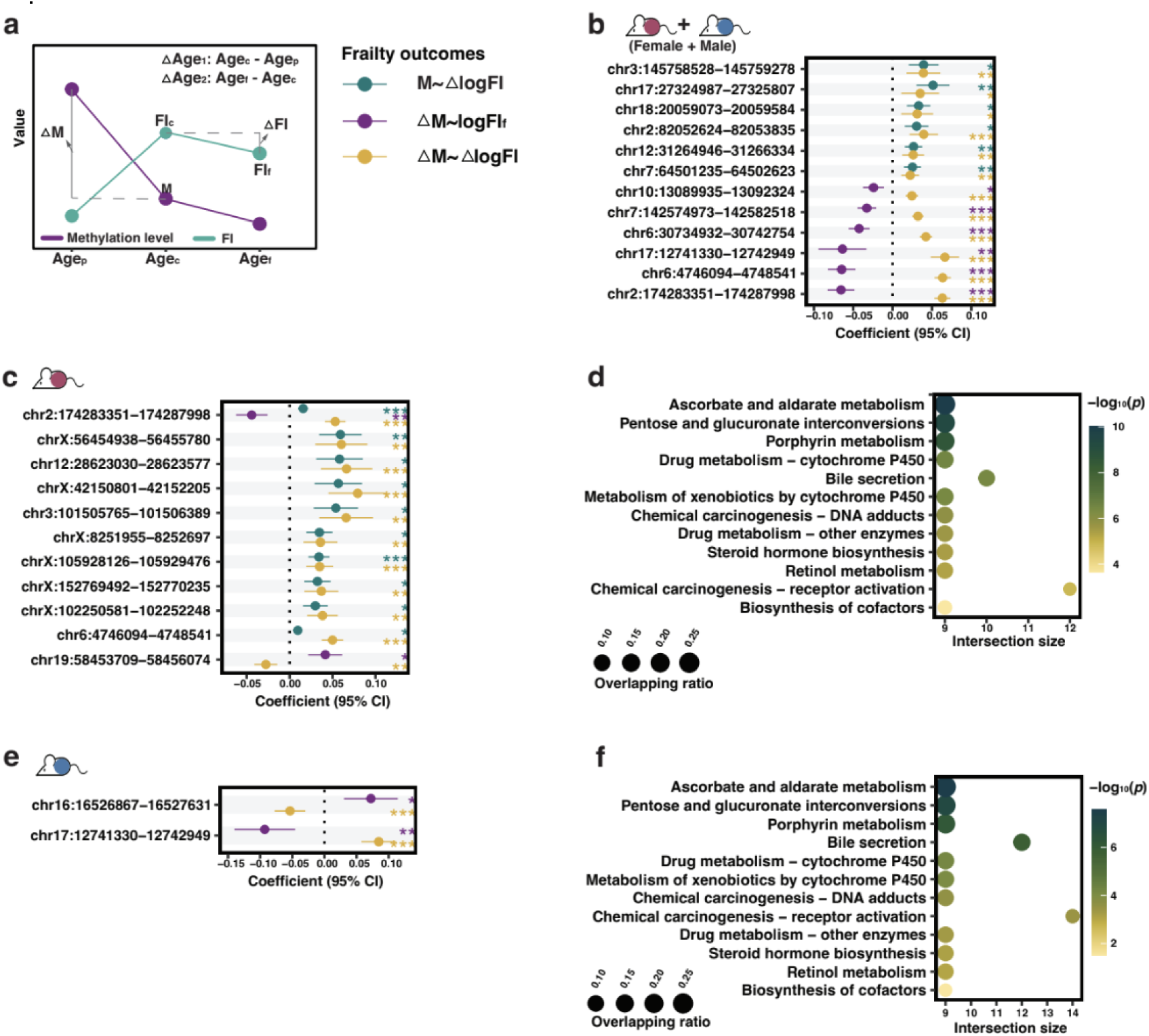
Association between differentially methylated regions and future frailty in the discovery cohort. **a.** Schematic diagram showing the dependent and independent variables in the association study of methylation M-value (M) and ΔM-value (ΔM, the difference in M-value between a prior age, Age_p_ and a current age, Age_c_; with adjustment for ΔAge_1_ being difference between Age_c_ and Age_p_) for each frailty-related differentially methylated region (DMR) with frailty outcomes, logFI_c_ (log-transformed Frailty Index at Age_c_), logFI_f_ (log-transformed FI at a future age, Age_f_; with adjustment for ΔAge_2_ being difference between Age_f_ and Age_c_), and □logFI (the difference between logFI_f_ and logFI_c_). For logFI_c_ and logFI_f_, when used as the outcomes to be associated with ΔM, further adjustment for logFI_p_ (log-transformed FI at a prior age) or logFI_c_ was performed. FI outcomes are distinguished by colors. **b.** Coefficients of 12 DMRs that present significance in the association with more than one FI outcome in the whole discovery cohort. The lines represent the 95% confidence interval of coefficients. **c.** Coefficients of 11 DMRs that present significance in the association with more than one FI outcome in females. **d.** Probe set enrichment analysis of pathways for CpGs within DMRs associated with future FI outcomes in females. **e.** Coefficients of 2 DMRs that present significance in the association with more than one FI outcome in males. **f.** Probe set enrichment analysis of pathways for CpGs within DMRs that present associations with future FI outcomes.

Within the whole cohort (males and females combined), we found no DMRs associated with future log-transformed FI (logFI_f_) but 14 DMRs associated with change of log-transformed FI (ΔlogFI). The majority (13/14) of identified DMRs associated with ΔlogFI displayed positive associations, indicating that more methylation in these regions was associated with higher frailty. We hypothesised that a change in methylation level over time (ΔM) may also be associated with or predictive of frailty, so tested the association of ΔM with two frailty outcomes (**Fig.5a**). We identified 7 DMRs whose change was associated with logFI_f_, and interestingly, these associations were all negative, implying that an increase in methylation at these regions over time was associated with lower frailty scores. Finally, we identified 170 DMRs whose change in methylation was associated with ΔlogFI. 153 of these presented positive associations, implying that a greater change in methylation at these regions was associated with a greater change in frailty. Overall these results indicate that temporal changes in methylation levels are predictive of subsequent frailty.

Assuming that significance of DMRs in more than one of these associations indicates sites of particular interest, we identified 12 DMRs that were significant across more than one of these analyses (**Fig.5b)**, and these DMRs are associated with ‘Amoebiasis’. Half of the DMRs showed the same direction of association for each model indicating that greater methylation (change) at these sites is associated with a greater increase in frailty over time. Notably, the other half DMRs exhibited associations in opposite directions across analyses, and the pattern persisted even after adjusting for the baseline FI (logFI_c_), suggesting complexity of these DMRs in associations with frailty level and progression (**Supplementary Fig.7a**). When considering the total 179 DMRs identified in the whole cohort as associated with our various frailty outcomes, we saw enrichment in genes such *Il27* and *Pou3f3*.

We completed the same analysis as above, stratified by sex. In females, 17 DMRs were positively correlated with ΔlogFI, but none with logFI_f_. For ΔM, 2 DMRs were associated with logFI_f_ and 142 with ΔlogFI. 11 DMRs were significant in more than one analysis (**Fig.5c**), including sites associated with *Gnasas1* and *Gnas*. The majority of these DMRs presented positive associations indicating that more methylation at these sites is associated with greater frailty in females, with 2 DMRs exhibited opposite directions, associated with *Gnasas1* and *Gnas*. The total 149 DMRs identified in these analyses for females were predominantly enriched for lipid metabolism pathways (**Fig.5d**). In males, no DMRs were associated with logFI_f_ or ΔlogFI, and there were 6 and 102 DMRs that showed significant associations between ΔM and logFI_f_ and ΔlogFI respectively for males. Of these, 2 DMRs were detected in more than one analysis (**Fig.5e**) and showed opposite directions, associated with the pathway ‘Virion-Herpesvirus’ and genes *Igf2r*. The total 106 DMRs identified in these analyses for males included CpGs linked to *Pantr1*. Intriguingly, only 12 DMRs were identified in both the female and male analyses, including CpGs associated with *Igf2r*.

Together we derived a total of 296 (32.0% of 925) DMRs after consolidating overlapping DMRs. The methylation levels or changes in levels of these DMRs were associated with subsequent changes in FI, serving as promising prognostic frailty biomarkers. These DMRs were enriched in lipid metabolism, cytochrome P450 and carbohydrate metabolism (**Fig.5f**). We also derived 22 DMRs that show significance in at least one association with frailty outcomes, and were linked to 118 genes, including *Gnas*, *Igf2r* and *Dsc2*. Overall this analysis identified several promising genomic regions for which methylation levels are associated with future frailty outcomes.

### Validation of FI associated DMRs

To determine whether the associations identified in the discovery cohort were conserved, we evaluated the relationship between the same DMRs and frailty outcomes in the validation cohort, which was treated with chronic NMN (an NAD precursor supplement^37^, see Methods). Specifically, we tested 1) associations between DMRs and FI, and 2) associations between DMRs and future FI outcomes, thereby validating the findings from the discovery cohort (**Supplementary Table 9**).

We estimated associations for the 925 frailty-related DMRs between the discovery and validation cohorts showed generally consistent directions of association (**Fig.6a**). A small number of DMRs (49, 5% of 925) exhibited reversed effect directions, however, these instances were almost exclusively limited to regions with coefficients near zero in the discovery cohort. This pattern suggests that direction flips primarily occur among DMRs with weak effects, consistent with variability expected from statistical noise rather than true biological discordance. In contrast, DMRs with larger absolute effect sizes demonstrated strong directional concordance, supporting the robustness of these associations across cohorts. In addition to analyses in the whole cohort, associations for sex-specific DMRs (771 for females and 397 for males) were evaluated within the corresponding male and female subgroups (**Fig.6b** and **c**). The overall patterns observed in the sex-stratified analyses were similar to those seen in the combined sample, with the strongest and most consistent effects replicated across cohorts.

**Fig.6.**
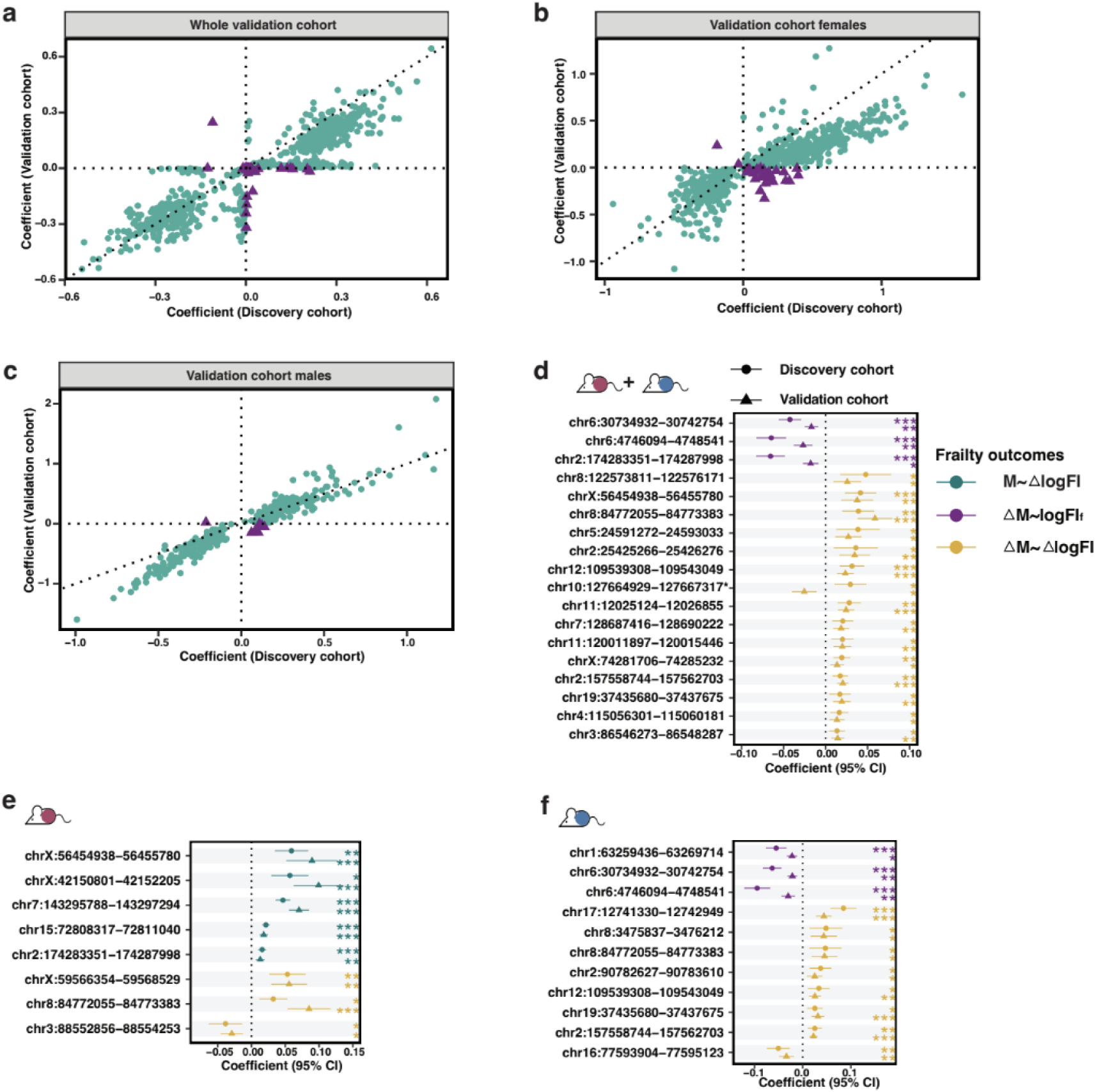
Validation of differentially methylated regions in the validation cohort. **a.** Comparisons of effect sizes of differentially methylated regions (DMRs) in association with frailty in the discovery and validation cohort in the whole cohort. Purple triangles represent those DMRs that show opposite directions of effect size between the discovery and validation cohorts. **b.** Comparisons of effect sizes of differentially methylated regions (DMRs) in association with frailty in females of discovery and validation cohort. **c.** Comparisons of effect sizes of differentially methylated regions (DMRs) in association with frailty in males of discovery and validation cohort. **d.** Coefficients of 18 DMRs that present consistent associations across the discovery and validation cohorts in the whole cohort analysis. The lines represent the 95% confidence interval of coefficients. FI outcomes are distinguished by colors. Circles represent the results from the discovery cohort and triangles for the validation cohort. **e.** Coefficients of 8 DMRs that present consistent associations across the discovery and validation cohorts in female-specific analysis. **f.** Coefficients of 11 DMRs that present consistent associations across the discovery and validation cohorts in male-specific analysis.

Considering future FI outcomes in both sexes, we found consistent associations for 18 DMRs with the same frailty outcomes as those found in the discovery cohort (**Fig.6d**), although the direction of association was flipped for 1 DMR, ‘chr10:127664929-127667317’ (**Supplementary Fig.7b**). This DMR is associated with *Stat6*, and may represent frailty-related sites affected by chronic NMN treatment. In females, 8 DMRs were consistently associated in the validation cohort, including 3 on the X-chromosome (**Fig.6e**). In males 11 DMRs were consistently associated across cohorts (**Fig.6f**), and none of the sex-specific associations exhibited a reversal in effect direction. These DMRs identified as showing consistent associations across cohorts represent strong candidates as markers of frailty, and include CpGs associated with *Gnas* and *Igf2r*.

### DNA methylation variability/entropy with frailty

Aging has been associated with more DNA methylation ‘noise’ or inter-individual variability^38^, but the association of DNA methylation variability with frailty is unknown. We investigated inter-individual variability across our identified frailty related CpGs either in the whole cohort, or in male and female subgroups by modeling both baseline methylation levels and individual-specific frailty effects using linear mixed models. Our analyses revealed noticeable variance in baseline methylation across mice, but minimal inter-individual differences in the trajectories (slopes) in frailty progression (**Supplementary Fig.8a**). This pattern suggests that while mice differ in their baseline methylation levels at fiCpGs, their rates of methylation change with frailty progression are relatively consistent (**Supplementary Fig.8b**). Examples are shown for cg37536990_TC11 and cg42178351_BC11, sites that showed the greatest variation in intercepts but least variation in slopes (**Supplementary Fig.8c**).

To identify specific CpGs that exhibit inter-individual variability in the association of methylation with frailty, we performed a Breusch-Pagan test to identify variably methylated probes (VMPs) that present intra-group variability. No VMPs were detected in females, or the whole cohort but we identified 135 VMPs (Benjamin-Hochberg adjusted *p* < 0.1) from male-specific fiCpGs. These male-specific VMPs were linked to genes such as *Esr1*, and were enriched in arrhythmogenic right ventricular cardiomyopathy related pathways. Furthermore, by calculating signal-to-noise (SNR) ratio, we determined 4 of these VMPs presented deterministic variations (SNR ratio > 10) (**Supplementary Fig.8d**) representing tightly regulated changes and associated with genes *Rag1* and *Rag2*, while 81 showed stochastic variations (SNR ratio < 1) (**Supplementary Fig.8e**), indicating random and unpredictable changes with the development of frailty, leading to genes *Esr1* and *Dsc2*.

To capture intra-individual (sample) epigenetic variation, we then calculated Shannon entropy^22^ (a measure of disorder or randomness of methylation patterns) across fiCpGs derived from our analysis. The results suggest a general increase in Shannon entropy, implying higher epigenetic instability, with the development of frailty (**Supplementary Fig.8f-h**), especially for age-dependent CpGs.

Taken together, our findings suggest that baseline DNA methylation levels vary significantly among individuals and that the progression of frailty is associated with increased epigenetic instability, and greater variability in methylation at specific sites, particularly for males.

### Epigenetic frailty clock

DNA methylation at specific CpG sites has been modelled to build epigenetic clocks that accurately predict chronological age in mice^39^. Here, we aimed to use methylation features to build an epigenetic clock that predicts frailty. To achieve this, we selected the top 100 DMRs (including 579 CpGs) that presented the greatest absolute coefficients with frailty (**Supplementary Fig.9**) and fit elastic net models in the discovery cohort to predict FI score. The first model ‘epigenetic frailty clock 1’ included age, sex and 68 CpGs (EFC1) and the second model ‘epigenetic frailty clock 2’ included sex and 105 CpGs (EFC2), excluding age, wherein 48 CpGs were shared and associated with genes including *Dsc2*. Within the discovery cohort, the epigenetic frailty clocks achieved adjusted *R*^2^ values at 0.59 (EFC1) and 0.60 (EFC2) (**Fig.7a**), with improved performance compared to baseline model (linear regression included age and sex) in both females and males. When applied to the validation cohort, performance in both males and females was similar to the baseline model (**Fig.7b**). When the estimated frailties from the models were plotted against time to death in both cohorts, they explained a larger proportion of variability than the conventional frailty index, especially in males (**Fig.7c**), and their performance was similar to or better than a simple age and sex model. The epigenetic frailty clocks display most value in estimating remaining lifespan for mice of the same age, where higher predicted frailties were associated with shorter lifespan expectancy, especially in older mice (**Supplementary Fig.10**). Despite the fact that there is clearly room for model improvement, these results suggest that frailty can be accurately predicted in aging mice using only methylation signatures.

**Fig.7.**
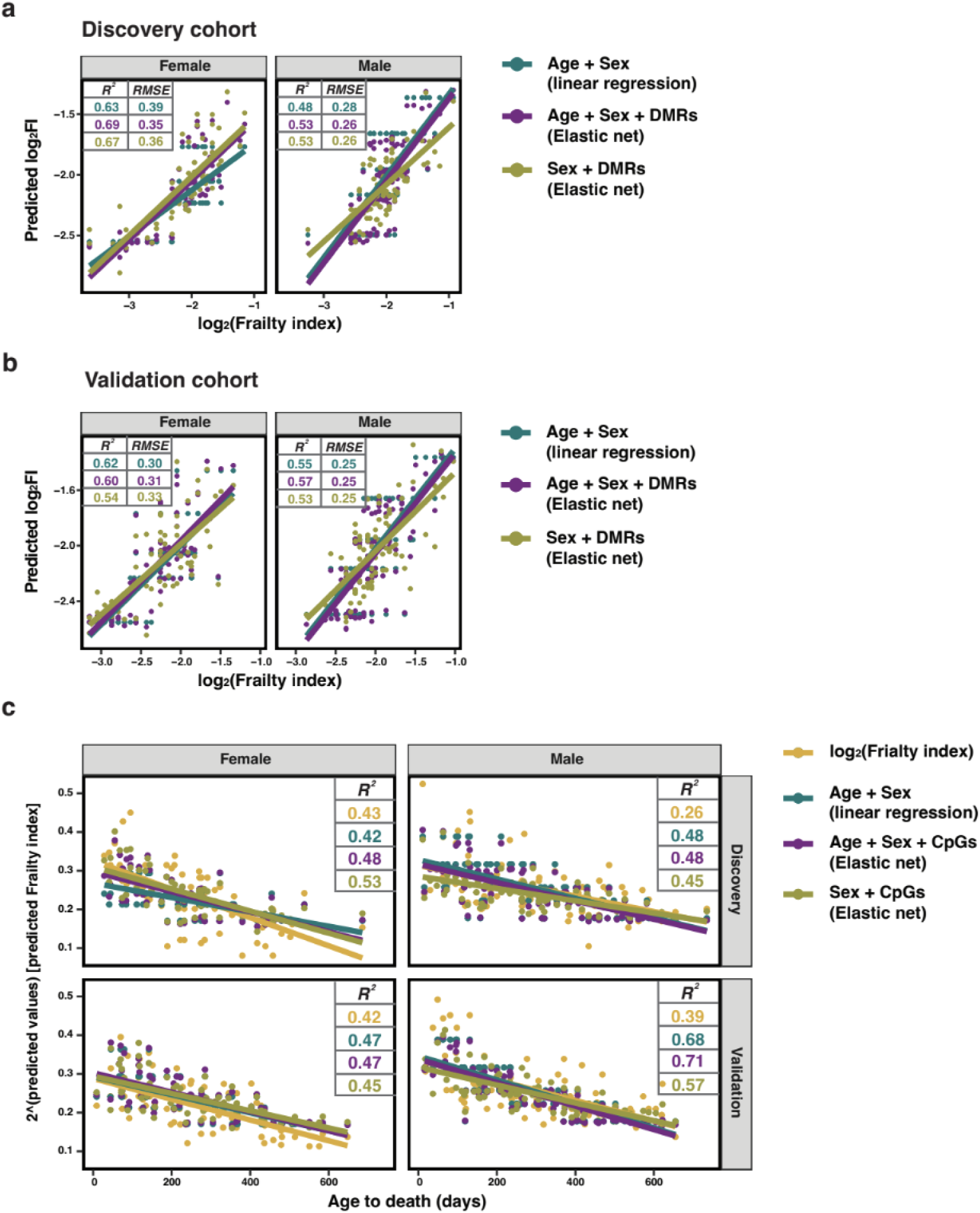
Performance of epigenetic frailty clock in the validation cohort. **a.** Comparison of the performance in female and male samples in the discovery cohort. Linear regression model (baseline model) was trained on age and sex. Two epigenetic frailty clocks were trained on sex, top 100 ranked DMRs (by absolute effect size association with frailty) and/or age. **b.** Model performance in the validation cohort stratified by sex. **c.** Proportion of data variability explained by log-transformed frailty index, age+sex model and epigenetic frailty clocks in female and male samples across the discovery and validation cohorts.

In summary, our analyses reveal DNA methylation signatures associated with components of frailty (AIFI and ADFI), sex dimorphisms and subsequent FI outcomes via a data-driven approach. A summary of representative epigenetic features is presented in **Fig.8**, highlighting the integrated methylation landscape associated with frailty progression.

**Fig.8.**
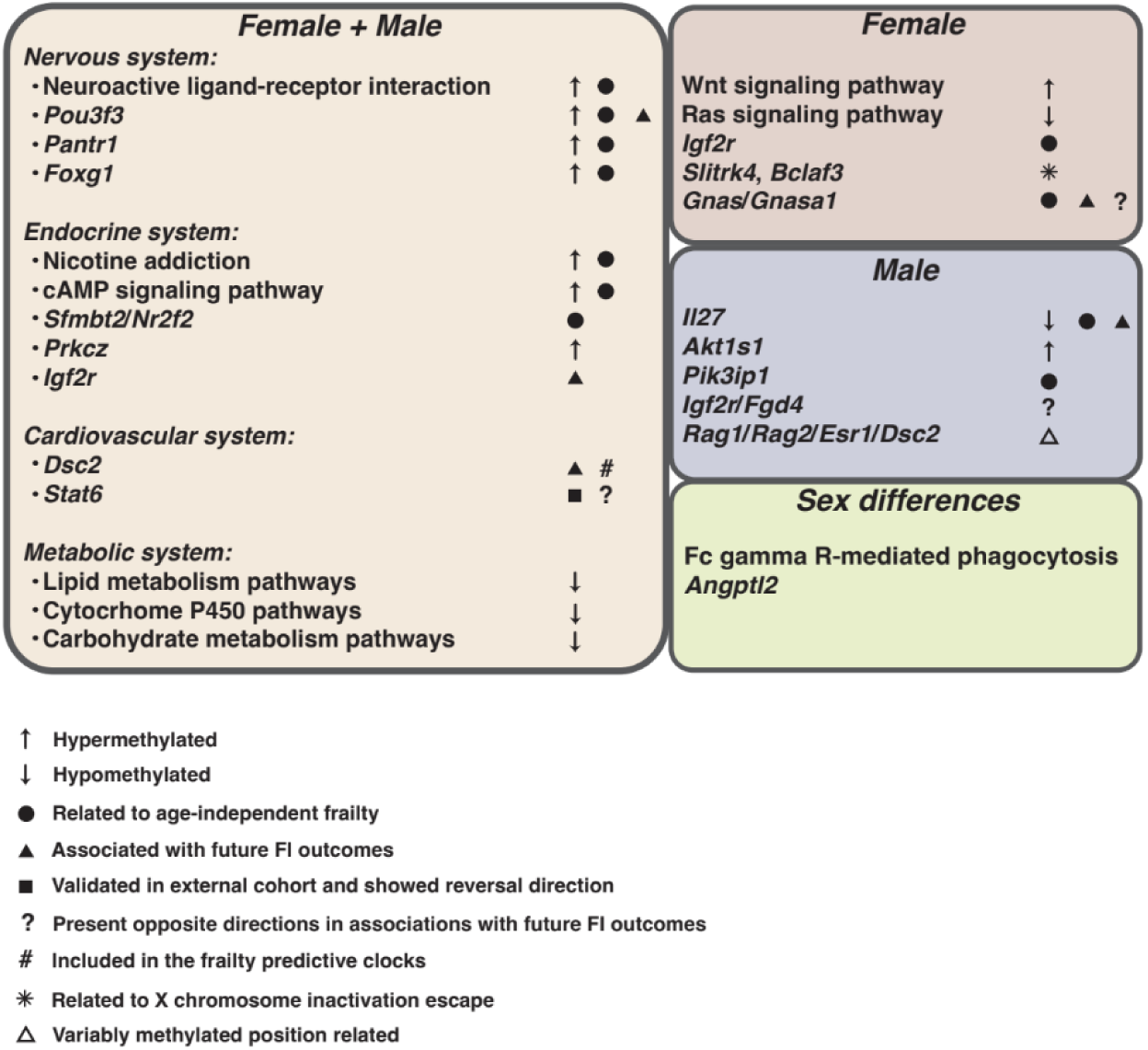
Frailty-related methylation signatures.

## Discussion

Using a longitudinal study of female and male mice, we identified both sex-independent and sex-specific methylation signatures of frailty. Overall, we found that age-independent frailty is linked with decreased methylation in genes related to nervous and endocrine system pathways, whilst age-dependent frailty is associated with increased methylation of genes related to signaling pathways. Female-specific methylation signatures suggest signaling, endocrine and nervous system related pathways that are distinct from those identified in males.

### Frailty related signatures independent of age

Markers that are associated with health in aging, rather than chronological age, may provide clues about underlying mechanisms of the aging process, rather than the passing of time. We sought to identify methylation features of frailty, a validated quantification of health in aging in both humans and mice. As frailty is strongly correlated with age, it was important that we identify methylation features of frailty, independent of age. Previous studies from us and others have explored age-independent metabolomic markers of frailty^40^, but only a few studies have explored DNA methylation frailty markers^25^. In this study, we employed a novel approach that integrates differential methylation analysis with linear mixed models to identify genomic regions whose methylation is associated with age-independent frailty (AIFI). Across the sex-inclusive subgroup (with adjustment for sex), we identified 412 genes (linked with 187 DMRs) whose methylation levels were associated with AIFI (**Supplementary Table2**), including *Pou3f3*, *Pantr1*, *Foxg1*, *Pantr2*, *Sfmbt2* and *Nr2f2*. These genes show either increasing or decreasing methylation levels with frailty, which are usually associated with decreased or increased gene expression respectively^41^, implying varied regulatory patterns for these genes in aging and frailty. For example, methylation levels at the *Foxg1* gene increase with frailty in our study, implying down-regulation of *Foxg1* expression in frailty. This observation aligns with a previous study demonstrating decreased mRNA and protein levels of *Foxg1* in progeria models of aging^42^. Long non-coding RNAs *Pantr1* and *Pantr2* are located adjacent to neurogenesis-related gene *Pou3f3*^43,44^, and *Pantr1* also participates in FOXG1 localization in both humans and mice^45^, which together are involved in neurodevelopmental disorders^46^. Given this, it is unsurprising that neuroactive ligand-receptor interaction pathways were enriched for these sites, suggesting that frailty is associated with Foxg1-linked disruption of neurodevelopmental regulatory pathways. *Sfmbt2* and *Nr2f2* (also known as COUP-TFII) are related to insulin and insulin-like growth factor 1^47,48^, which are also linked to insulin resistance^49^. Frailty is associated with increased insulin resistance^50^, and our results suggest that this metabolic alteration may be mediated by hypermethylation of genes within nicotine addiction pathway, resembling the effects of chronic nicotine exposure^51^. Overall, our results suggest that the established link between frailty and changes in nervous and endocrine systems^50,52,53^, may have underlying epigenetic origins. The genes and pathways identified from our methylation analysis are particularly interesting for further analysis as underlying markers of health in aging, independent of chronological age.

### Frailty-related methylation features

Investigating methylation regions contributing to both frailty and aging can also provide insight into common genes and pathways underlying the biological mechanisms for age and frailty. Here we generated a joint set of 925 frailty-related DMRs, that were both positively and negatively associated with frailty and enriched for lipid metabolism and cytochrome P450 related pathways. Although no consistent pattern was observed across all DMRs, our results suggest, in general, hypomethylation of genes involved in lipid metabolism and cytochrome P450, implying increased expression of these genes with frailty and aging (**Fig.4g**). Given the complexity of these pathways and their numerous components, the precise mechanisms, particularly those involving hypomethylation, underlying frailty progression remain to be fully elucidated. Some specific examples can be highlighted however, such as the gene *Prkcz,* the methylation of which was positively associated with frailty. Previous work has shown hypermethylation of *Prkcz* is linked to type II diabetes mellitus^54^ which is associated with frailty in humans. Additionally, we observed hypermethylation of genes involved in cAMP and Wnt signaling pathways with frailty (**Fig.4f**), both of which have been implicated in aging by previous studies^55,56^, indicating loss of cell function in frailty progression. Together, the identified methylation patterns suggest both gene hypermethylation and hypomethylation may contribute to frailty development.

Additionally, we applied linear mixed models to frailty-related DMRs to look at their specific univariate association with subsequent frailty outcomes. The longitudinal nature of our data allowed us to complete the first study looking at the association of both changes in methylation levels at DMRs over time, and changes in frailty over time. Although we didn’t identify any regions for which methylation levels predicted future frailty, we did find 296 DMRs (and changes in DMR methylation, ΔM) that were associated with a change in frailty (ΔlogFI). These were enriched, especially in females, in lipid metabolism, cytochrome P450 and carbohydrate metabolism (**Fig.5d**), identified in both females and males from frailty-related DMRs. Epigenetic regulation of these pathways may play an important role in the development of frailty, and these sites represent exciting clinically relevant targets for follow up as potential biomarkers to predict those at risk of becoming more frail in the near future.

We also found 22 DMRs that were significant in at least two of our longitudinal association analyses, suggesting they may be of particular value as markers of frailty. These informative DMRs are associated with genes involved in the endocrine system (*Gnas*/*Gnasas1* in females and *Igf2r* in males), suggesting pathways within the system may represent core biological mechanisms with frailty progression. Intriguingly, the majority of DMRs showed consistent directions in their associations with frailty outcomes, nevertheless 7 (< 1% of 925) DMRs presented inconsistent directions, that is, the methylation change of DMRs presented opposite directions in associations with future FI and FI change. We performed a sensitivity test, adjusting for baseline FI, to rule out its confounding effect, however, opposite directions persisted, likely reflecting stochastic variation or biological and environmental context-dependent methylation-frailty relationships. From these DMRs, we identified *Gnasas1* and *Gnas* that are linked to cAMP signaling pathway^56^ in females, as well as *Igf2r* in males. All three genes point towards the endocrine system, indicating the complexity of its involvement in the epigenetic regulation of frailty.

Importantly, and often ignored in other frailty biomarkers studies, we confirmed that many of the same DMRs were associated with frailty outcomes in an independent validation cohort. Our validation cohort was part of a larger study that received long-term NMN treatment^37^. 28 DMRs were validated in this external cohort, such that the associations with frailty outcomes persisted across studies. These DMRs again involved sites associated with *Gnas* and *Igf2r*, suggesting the endocrine system. While the majority of DMRs presented consistent directions of associations with future FI outcomes across the discovery and validation cohorts, the DMR associated with *Stat6* exhibited an opposite trend, shifting from a positive association in naturally aging mice to a negative association in mice with NMN treatment. Given that inhibition of *STAT6* has been shown to mitigate muscle loss and improve muscle performance^57^, this reversal may reflect biological modulation of the STAT6 signaling pathway, potentially influenced by NMN treatment. These results suggest that associations between the majority of methylation features with frailty outcomes may be universal even under interventions, but a few DMRs of which the associations were altered by the intervention. Therefore, these features should be investigated further.

### Sex differences in frailty related methylation features

The sex-frailty paradox has been extensively investigated, with evidence indicating that women tolerate the deleterious effects of frailty better than men^58^. Therefore, elucidating underlying biological mechanisms contributing to sex differences in frailty is essential for advancing our understanding of frailty and addressing its clinical implications.

X chromosome inactivation is a process to balance gene dosage in females. While the majority of genomic X regions follow this pattern, some regions escape from this regulation and remain transcriptionally active and have been reported to be functional in aging^59^. By comparing beta values of CpGs within the X chromosome in both sexes, we were able to detect 13 X chromosome inactivation escape (XCA)-related CpGs associated with frailty. These regions were associated with the genes *Slitrk4* and *Bclaf3,* suggesting a female specific role for their regulation in frailty development, although no established links between these genes and frailty have been reported to date. Nonetheless, their escape from XCI in blood cells highlights the need for further investigation into their potential regulatory roles.

Beyond the X chromosome, female specific DMRs associated with frailty were linked to 66 genes, including *Igf2r*. *Igf2r* encodes the insulin-like growth factor 2 (IGF2) receptor that binds IGF2, which is implicated in at least two of the hallmarks of aging, cellular senescence^60^ and mitochondrial dysfunction^61^. Our results confirm the involvement of these markers in aging and suggest a possible link to frailty, particularly in females. Additionally, regarding enriched pathways for female-specific DMRs (**Fig.4c**), Wnt signaling and neuronal pathways (circadian entrainment^62^, Ras signaling pathways^63^ and retrograde endocannabinoid signaling^64^) have been linked to frailty previously, although not in a sex specific manner.

In males, specific genes whose methylation was positively and negatively associated with frailty include *Akt1s1* and *Il27* respectively. *Akt1s1* serves as a regulator for the mTORC1 signaling pathway^65^ with growing interest in its role in frailty^66^. *Il27* is associated with inflammation^67^ known to play a significant role in frailty^68^. While *Il27* has previously been linked to frailty in a longitudinal study, this association was observed in women^69^. These results suggest different mechanisms involved in the development of frailty in females and males, reflected by distinct CpG-enriched gene markers and pathways with established ties to frailty. This highlights the need for further studies to explore the potential roles of these markers in frailty, especially in a sex-specific context.

In addition to identified sex-specific differentially methylated regions, we also identified several shared regions between males and females that exhibited distinct sex differences. These included regions such as those associated with gene *Angptl2*, which showed an opposite direction of association with frailty in males and females, and has been previously shown to regulate energy metabolism in mice in a sex-dimorphic capacity^70^. We also identified 36 DMRs which were associated with age-independent frailty in males, but were classified as age dependent in females. These regions were enriched in Fc gamma R-mediated phagocytosis, which has been reported to exhibit contrasting trends in activity during aging between sexes^71^. These findings highlight the need for further investigation into the sex-specific regulation of these pathways in the context of frailty.

There is growing awareness that aging is associated with not just predictable changes in methylation at specific genomic sites, but also a general increase in variability of methylation both within and across samples^21^. Here, our analysis, focused on CpGs within frailty-related DMRs, revealed substantial inter-individual variability in baseline DNA methylation levels across frailty-related CpGs, yet relatively consistent trajectories of methylation change with frailty progression. Notably, sex-specific patterns emerged, with male-specific fiCpGs showing greater intra-group variability. We identified 135 variably methylated probes (VMPs) from male mice, linked to genes such as *Esr1* and enriched in cardiomyopathy-related pathways. *Esr1*, which encodes the estrogen receptor and has been implicated in frailty and aging processes in females^72^, was unexpectedly identified among male-specific VMPs, suggesting a possible role in its regulation in the development of males also. Furthermore, signal-to-noise analysis indicated both deterministic and stochastic methylation changes in males, while Shannon entropy increased with frailty in both sexes, suggesting rising epigenetic instability. These findings highlight sex differences in methylation variability and suggest that epigenetic responses to frailty may be more heterogeneous in males.

Together, these findings reveal several distinct layers of sex differences in frailty: the contribution of X chromosome inactivation escape for females, differential methylation signatures and pathways across sexes, sex-divergent regulation of shared regions, and greater epigenetic variability in males. This multi-dimensional evidence underscores the importance of incorporating sex-specific analyses in studies of frailty and suggests that tailored interventions may be needed to address the unique biological mechanisms contributing to frailty in females and males.

### Epigenetic frailty clock

Epigenetic clocks are widely used in aging studies to predict biological age in humans^10,12,73^ and mice^74^, and to investigate associations between aging and health-related outcomes^75,76^. Quite recently, there is a growing focus on building models to predict frailty^25,40^, underscoring the potential of DNA methylation markers to predict frailty. For example, a recent study derived and validated an epigenetic frailty risk score (eFRS) using CpG sites associated with a frailty index in older adults, demonstrating robust associations with frailty across independent population-based cohorts^25,40^. Here, we build a clock to directly predict frailty in mice from methylation features. Our model provides robust prediction of frailty in our discovery cohort (**Fig.8a**) and performance is similar to that of an age+sex only model in females for the validation cohort, but outperforms the simple model in males (**Fig.8b**). These results provide preliminary evidence that it is possible to predict frailty in mice using DNA methylation data but suggest further work should be done in large datasets to develop a more universal frailty clock.

There are some limitations to this study. Our validation dataset was relatively small, and mice were treated with NMN that may alter DNA methylation levels and other outcomes. We suggest future work should validate these potential frailty markers in larger cohorts, as well as in other mouse strains and humans. Additionally, the sample size for methylation data is relatively small, especially at the older ages, which might decrease the power of statistical analysis. Survival bias is also an issue to consider, as mice died over the course of the study and only those that were longest lived made it to timepoint 4 and 5. For future work, it will be ideal to conduct studies in a broader age range with an increased number of mice.

In summary, we performed a longitudinal study of naturally aging female and male mice looking at DNA methylation of frailty with a focus on age-independent frailty. We found nervous and endocrine system pathways are linked to age-independent frailty and highlighted sex dimorphism in the associations between DMRs and frailty.

## Methods

### Mice samples

Mice used in this study are from a larger intervention study, so detailed methods can be found in the previous manuscript^37^. Briefly, C57BL/6JNIA mice, female (*n* = 40) and male (*n* = 49) were obtained from the National Institute on Aging (NIA) Aging Rodent Colony, among which, 20 female and 24 male mice were subjected to nicotinamide mononucleotide (NMN) treatment. Mice were group housed (4-5 mice per cage, although over the period of the experiment mice died and mice were left singly housed), at Harvard Medical School in ventilated microisolator cages, with a 12-hour light cycle, at 71°F with 45-50% humidity. Mice were fed AIN-93G Purified Rodent Diet (Dyets Inc, PA). All animal experiments were approved by the Institutional Animal Care and Use Committee of the Harvard Medical Area. In order to investigate aging and frailty related DNAm level changes and mechanisms in naturally aging mice, we used non-NMN treated mice (females, *n* = 20; males, *n* = 25) as the discovery cohort for principal component analysis, differential methylation analysis, sex stratified analysis, association study, and DNAm frailty clock model building (**Fig.1**). We then tested the selected CpG sites (CpGs) features and model in the NMN treated mice (validation cohort).

### Mouse Frailty assessment

Behavioral and clinical variables for clinical frailty index were measured in both the discovery and validation cohorts, at each time point (**Supplementary Table 1**). We utilized the mouse clinical frailty index^2^ (FI) that contains 31 health-related items for this study. Briefly, mice were scored either 0, 0.5 or 1 for the degree of deficit they showed in each item with 0 representing no deficit, 0.5 representing a mild deficit and 1 representing a severe deficit.

### Blood collection and processing

Mice were fasted for 5-6 hours, anesthetized with isoflurane (5%) and then blood was collected from the submandibular vein with a lancet (maximum 10% of mouse body weight, approx. 200-300 μl), into a tube containing 20 μl of 0.5M EDTA. Blood was mixed and stored on ice. Whole blood was centrifuged at 1500×g for 15 mins, plasma was removed and PBMCs frozen at -80°C for subsequent DNA extraction.

### DNA extraction and DNA methylation array analysis

PBMC DNA extraction was performed as previously described^77^. In brief, RBC lysis buffer was used to remove red blood cells (155 mM NH4Cl, 12 mM NaHCO3, 0.1 mM EDTA, pH 7.3) then proteinase K (20mg/ml) in TESR buffer (10mM Tris-HCl, 25 mM EDTA, 0.5% SDS, 20ug/mL RNAse) was added to the pellet, and incubated at 65°C overnight. DNA was isolated using SPRI beads and eluted in 10 mM Tris-HCl pH8. Qubit (ThermoFisher) was used for DNA QC, and samples were then processed by FOXO Technologies using the Infinium Mouse Methylation BeadChip (Illumina, CA, USA).

### DNA methylation data processing

Raw IDAT files from the DNA methylation array were subjected to the SeSAMe^78,79^ pipeline for data preprocessing and quality control using code ‘TQCDPB’. Specifically, the pooBAH algorithms were included in the pipeline to mitigate the batch effect^80^. The numbers of DNA methylation array data at each time point are summarized in **Supplementary Table 1**. We excluded non-CpG sites and Single Nucleotide Polymorphism probes, filtered out CpG sites that were detected in less than 5% of samples in the discovery cohort or presented near-zero variance (failing detection in more than 5%), and obtained 242,599 CpGs. We then converted the beta-value derived from the pipeline to M-value for the ensuing analysis^81^. All statistical analyses were based on M-values if not stated otherwise. The genomic analysis presented in the study was performed using the mm10 mouse genome reference build, as described in the Illumina manifest file associated with the Infinium Mouse Methylation BeadChip.

### Selection of frailty-related DMRs

The whole process of DMR selection was performed respectively in the sex-inclusive subgroup (including both female and male samples but excluding male samples at the T5 time point) and sex-specific subgroups (female and male samples separately). DNA methylation M-value data were subjected to Epigenome-Wide Association Study (EWAS) analysis by using the ‘limma’ pipeline^82^ to associate DNA methylation levels (M-value) with log-transformed FI. Design matrices included sex and sex by log-transformed frailty interaction term (sex factor excluded in sex stratified analysis), without assigning a reference level^83^. The data along with the multi-factor design matrix were then subjected to linear modeling with the intra-block correlation based block on mouse ID, and empirical bayes smoothing of standard deviations. We then selected differentially methylated positions (DMPs) by controlling for a 5% Benjamini-Hochberg (BH) false discovery rate (adjusted *p*-values < 0.05). DMPs were subjected to ‘DMRcate’ pipeline^84^ to identify potential biological functional DMRs. Gaussian kernel bandwidth for smoothed-function estimation (*lambda =* 1000) and Scaling factor for bandwidth (*C =* 2) were set up according to the recommendation of the package for array data. We used either absolute t-statistic or square root of F-statistic as weight. DMRs with at least 5 CpGs were determined by controlling the threshold of Stouffer summary transformation of the individual CpG *p*-values at 0.05.

Frailty is highly related to age^2^, hence we are interested in DMR clusters that are related to the age-independent component of frailty (AIFI) as well as the age-dependent component of frailty (ADFI). Therefore, we applied an additional selection step at DMR levels to associate DMR with both components of frailty. DMRs are regarded as possible functional regions^85^, hence we included all CpGs within DMRs (i.e. DMR-related CpGs including both DMPs and non-DMPs). We followed the *coMethDMR* workflow^35^. Briefly, we excluded CpGs that present low co-methylation levels within the DMR by setting up a threshold for ‘*rdrop’* statistic (correlation between each CpG with the sum of methylation levels in all other CpGs) at 0.4. In the sex-inclusive analysis, methylation level was adjusted to exclude sex factor prior to the correlation, and was subjected to the ensuing analysis. Methylation level in sex-specific analysis remained unadjusted. For each remaining DMR (no. of CpGs ≥ 5), we applied linear mixed models using methylation levels as the dependent variable and either log-transformed FI (with and without adjustment of age) or age as the independent variable, allowing random mouse individual effect and random CpG effect. Associations between DMRs and frailty or age were determined by controlling for a 5% BH false discovery rate (adjusted *p*-values < 0.05). Wald test of coefficient was used to assess the association between the DMR and frailty or age. DMRs that only presented significance in association with logFI with adjustment for age (i.e. AIFI) were classified as age-independent DMRs (aiDMRs); those only presented significance in association with age (i.e. ADFI) were classified as age-dependent DMRs (adDMRs); and those presented significance in associations with both AIFI and ADFI were classified as dualDMRs.

### X chromosome inactivation identification

We followed a decision tree workflow^36^ to identify the status of X chromosome inactivation (XCI), analyzing a total of 10,338 CpG sites across all female and male samples. Briefly, for each CpG site, average methylation levels were calculated for males and females respectively, and were then classified according to methylation levels and differences between sexes. CpG sites were predicted to escape XCI when both male and female averages were unmethylated (< 0.15% methylation) and exhibited similar methylation ranges (i.e., overlapping ranges or < 10% difference between male and female averages); predicted to be subject to XCI when males and females displayed distinct methylation ranges with a difference in average methylation > 10%; predicted as variably escaping XCI when exhibiting a >10% difference in average methylation but overlapping methylation ranges; and considered unclassifiable otherwise.

### Statistical analysis

For principal component (PC) analysis, 153 samples within the discovery cohort, each with M-value for 242,599 eligible CpG sites, were included. We derived PC1 to PC15 and for each PC as the dependent variable, we applied linear regression models and obtained *p*-values, whereby each of the following factors, mouse ID, time points, sex, cage (categorical variables), and age at assessment (continuous variables), was used as the independent variable, which were then clustered by *p*-values according to Euclidean distance. CpG-associated genes and KEGG pathways were selected by using R package ‘KnowYourCG’ to address data sparsity in array-based datasets^86^. Although the analysis focused on the DMR level, individual CpGs within those regions were used for enrichment due to the sparse coverage of the methylation array. The final set of frailty-related DMRs were associated with log-transformed frailty index by applying linear mixed models adjusting for sex. *P*-values for DMR corrected by BH procedure were derived for Manhattan plot visualization. Within the association study, for each DMR in the final set, we applied linear mixed models by using 1) logFI_c_, log-transformed FI at current age (Age_c_), 2) logFI_f_, log-transformed FI at a future age (Age_f_), and 3) log-transformed FI changes (logFI_f_ - logFI_c_, ΔlogFI) respectively, as the outcome and M-value of methylation level or the difference in M-value between a past age (Age_p_) and Age_c_ (ΔM-value) as the independent variable adjusted for age and sex. We then selected DMRs presenting significant association by controlling for a 5% Benjamini-Hochberg (BH) false discovery rate (adjusted *p*-values < 0.05).

## Data availability

Mice metadata, DNA methylation data, and R markdown file for data analysis are available at https://github.com/Kane-Lab-ISB/Longitudinal-DNA-methylation-analysis-in-mice. The raw datasets will be deposited in the Gene Expression Omnibus (GEO) and released publicly upon publication of the manuscript. All data will be made available to editors and reviewers during peer review.

## Author contributions

A.E.K., P.G and D.A.S. conceived and designed the study. A.E.K., M.M and P.G. performed the experiments. D.Z. conducted the data analysis, with contribution from P.G. and S.F.. D.Z., S.F., and A.E.K. drafted and revised the manuscript with help from all authors. All authors have read and agreed to the published version of the manuscript.

## Correspondence

Correspondence to Alice E. Kane.

---

**Supplementary Fig.1.**
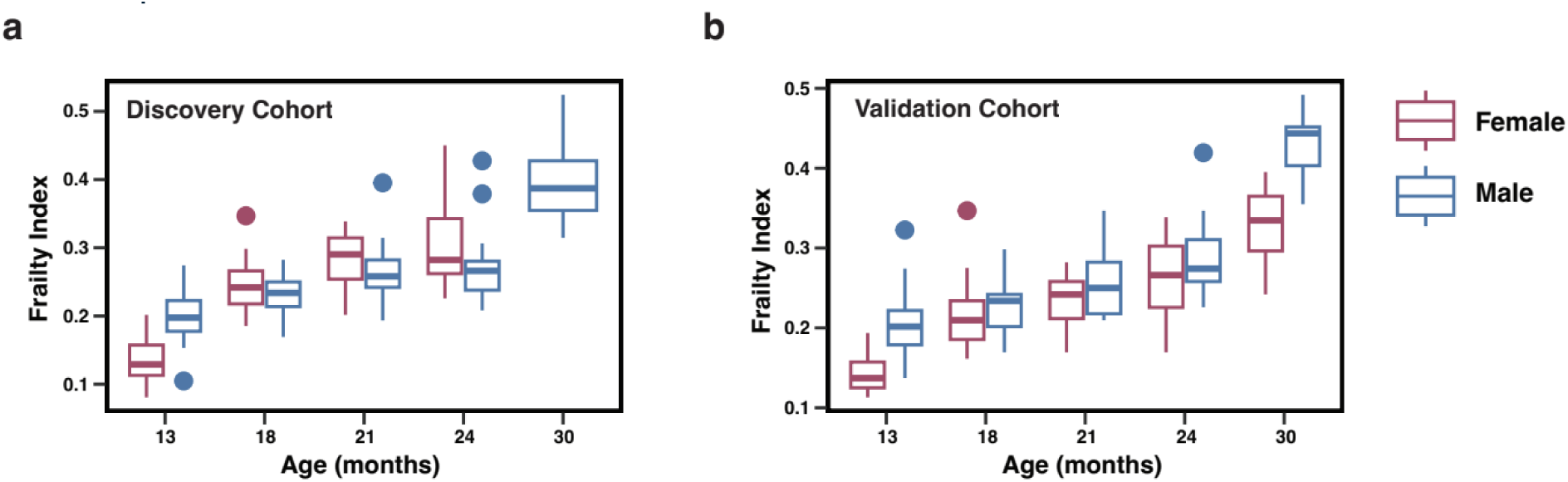
Distribution of frailty index. Frailty index (FI) for each mouse at each time point was derived from the 31-Item Clinical Frailty Index test. **a.** Boxplot of FI in the discovery cohort stratified by sex and time points (age in months). **b.** Boxplot of FI in the validation cohort stratified by sex and time points (age in months). The box represents the interquartile range from the 25th to the 75th percentile, the line inside the box indicates the median (50th percentile). Outliers (beyond 1.5 times the interquartile range) are plotted as individual points.

**Supplementary Fig.2.**
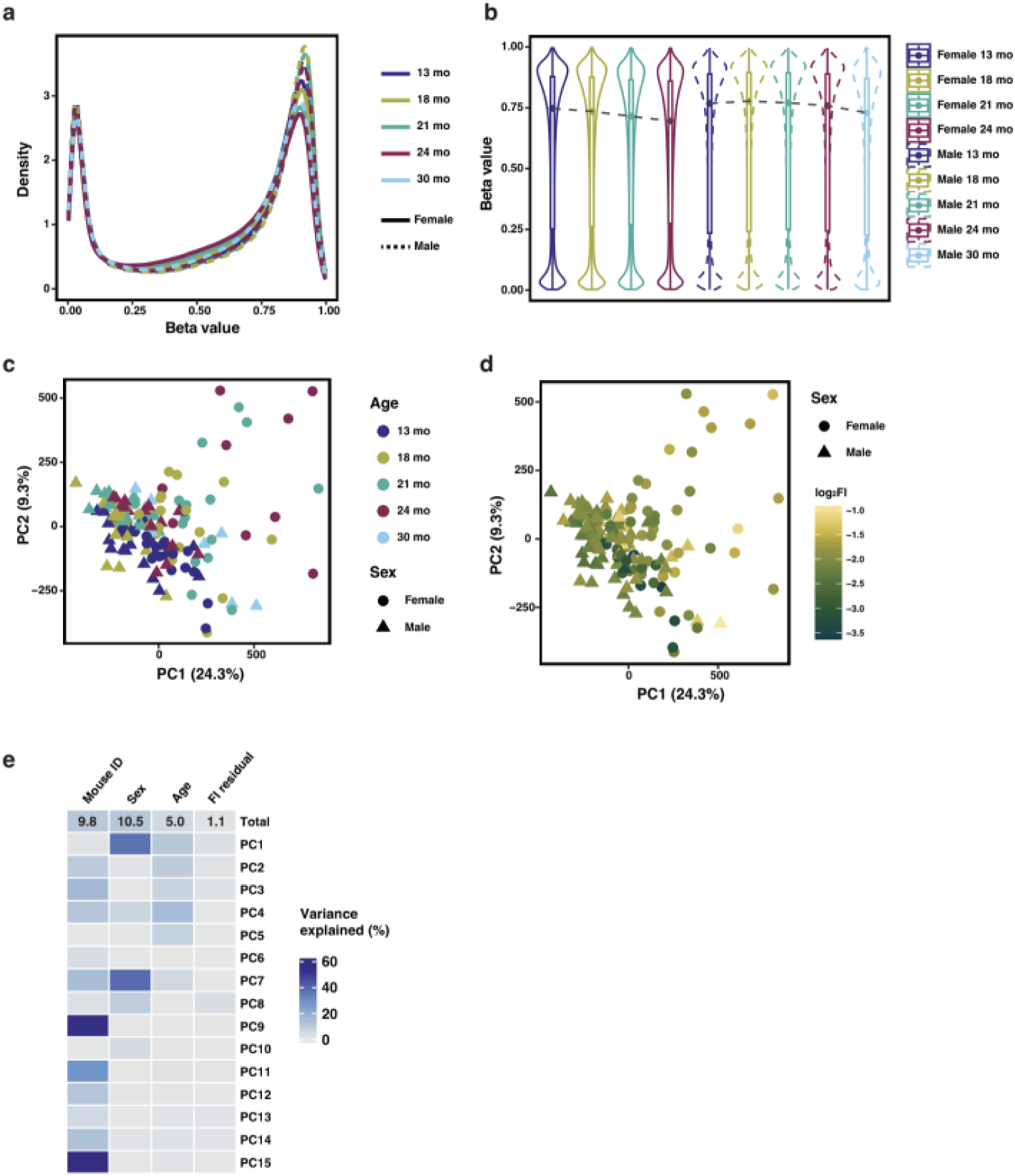
Global DNA methylation changes across time and sex. DNA from peripheral blood mononuclear cell samples in the discovery cohort (*n* = 153) from 20 female and 25 male mice was analyzed using a DNA methylation array. After quality control and filtering, 242,599 CpG sites were retained for downstream analysis. **a.** Distribution of beta values by density plot stratified by sex and age (13 mo, 18 mo, 21 mo, 24 mo, and 30 mo). **b.** Distribution of beta values by violin and boxplot stratified by sex and age. Dashed lines connect the median beta values across time points to illustrate temporal trends in global methylation levels. **c.** Principal Component analysis (PCA) plot (PC1 vs. PC2) showing sample variation by sex and time point. Analysis was performed on M-values (logit-transformed beta values). **d.** PCA plot (PC1 vs. PC2) showing potential associations between methylation patterns and frailty (log-transformed frailty index, log_2_(FI)). **e.** Heatmap showing total sample variance and PC variance (%) explained by mouse ID, sex, age and FI residual (age effects regressed out). Variance partitioning analysis was performed using linear mixed models to estimate the variance attributable to variables.

**Supplementary Fig.3.**
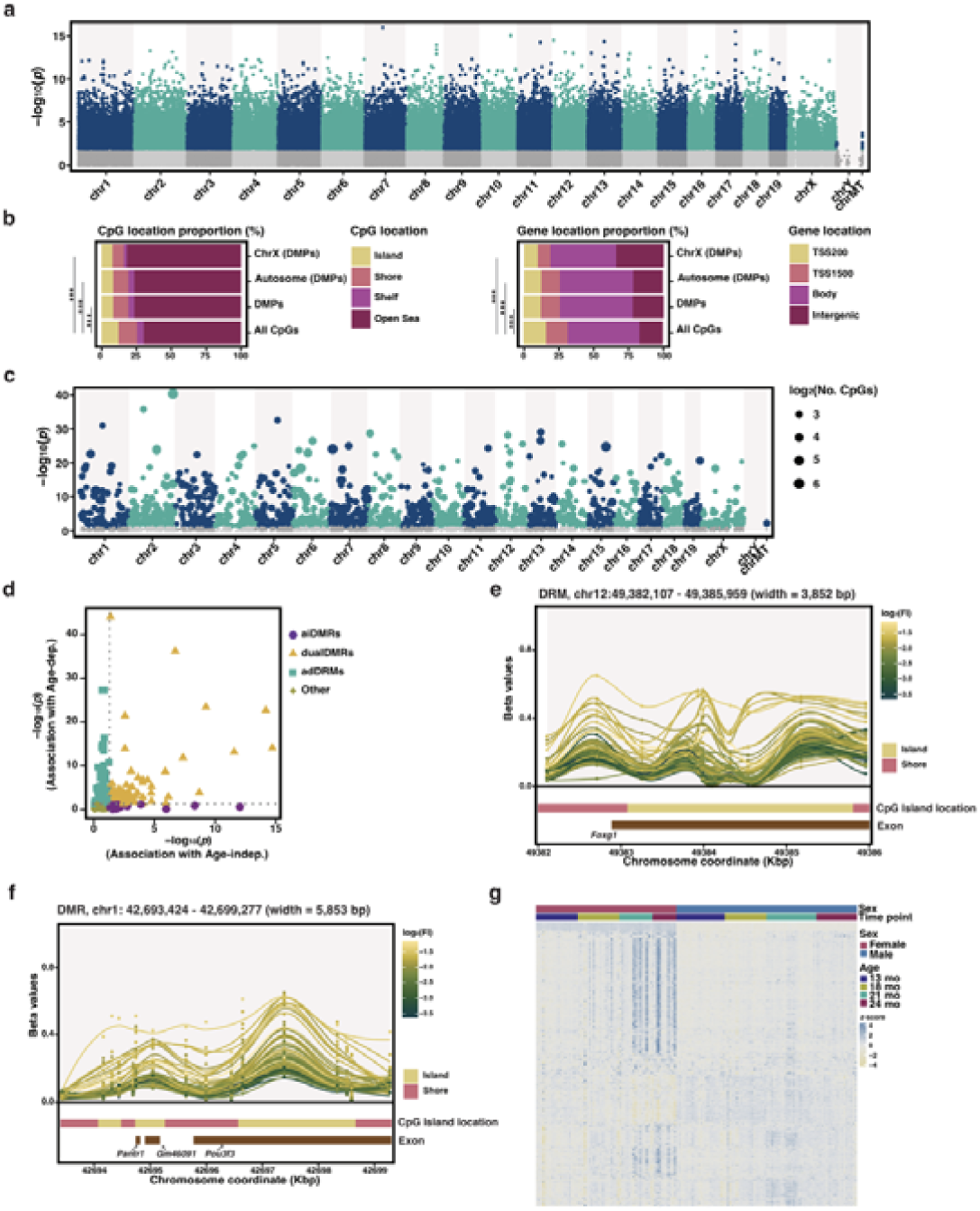
Visualization and characterization of frailty-related DMPs and DMRs in sex-inclusive subgroup. **a.** Manhattan plot showing the differentially methylated positions (DMPs) associated with frailty. DMPs were selected by controlling for a 5% Benjamini-Hochberg (BH) false discovery rate. **b.** Barplot showing the proportion of CpG and gene location categories. Proportions of ‘Island’ and ‘TSS200’ among all CpGs included in the analysis were compared to those of DMPs in all chromosomes, autosomes and X chromosomes by chi-square test, with *** for *p* < 0.001. **c.** Manhattan plot showing the differentially methylated regions (DMRs) associated with frailty. DMRs were identified by Stouffer’s Z-score method and selected based on a *p*-value threshold of 0.05. The size of each point corresponds to the number of CpGs included. **d.** Scatter plot showing *p*-values of DMR in associations with age-independent (associated with frailty adjusting for age) and age-dependent frailty. Each point represents a DMR, and is colored according to classification. **e.** Heatmap showing association between modules with frailty and age. Module eigenvalues were subjected to association with log-transformed FI (adjusted for sex or adjusted for sex and age) and age by applying linear mixed models. Significance was determined by *p*-values at a cutoff of 0.05, with * for *p* < 0.05, ** for *p* < 0.01, and, *** for *p* < 0.001. **f** and **g.** Illustrative examples of differentially methylated regions. Each point represents the methylation beta value of a CpG within the genomic location and CpGs from the same sample are connected by smoothed lines fitted using the LOESS method. Both points and lines are colored according to the level of log-transformed frailty index (log_2_FI). Distribution of CpG location categories (colored according to the category) and exons is presented in panels.

**Supplementary Fig.4.**
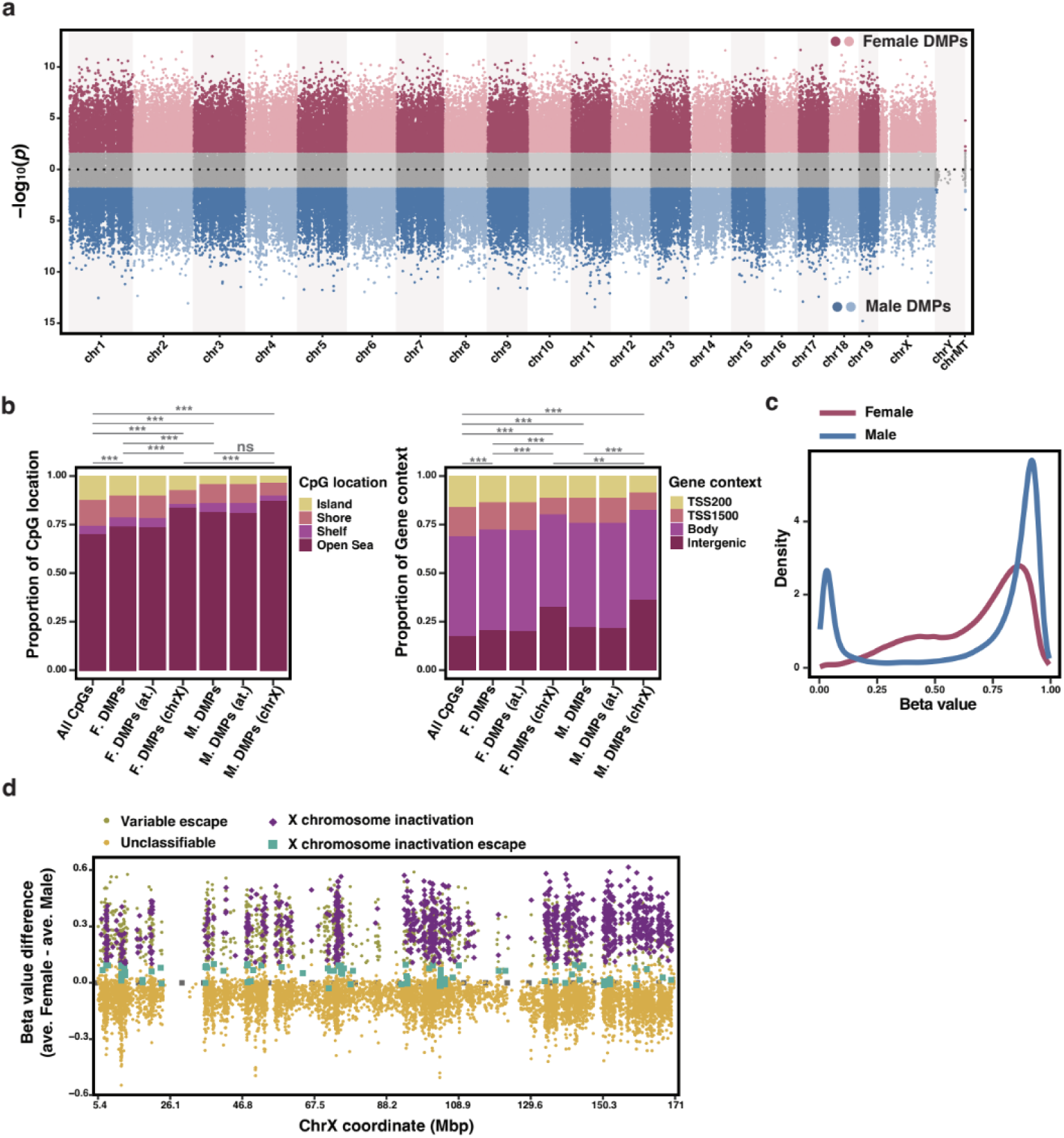
Selection of CpGs linked to X chromosome inactivation and escape. **a.** Manhattan plot showing the differentially methylated positions (DMPs) associated with frailty in females and males. DMPs were selected by controlling for a 5% Benjamini-Hochberg (BH) false discovery rate. **b.** Barplot showing the proportion of CpG and gene location categories in females (F.) and males (M.). Comparisons of proportions of ‘Island’ and ‘TSS200’ among all CpGs included in the analysis (All CpGs), DMPs in all chromosomes, autosomes (at.) and X chromosomes (chrX) were performed by chi-square test, with ** for *p* < 0.01 and *** for *p* < 0.001. **c.** Distribution of beta values of X chromosome linked CpGs in females and males. **d.** Identification of CpGs types related to X chromosome inactivation. CpGs were classified by beta value difference and overlap of CpG methylation ranges between females and males. CpG types are colored and shaped according to the classification (escape, X chromosome inactivation escape).

**Supplementary Fig.5.**
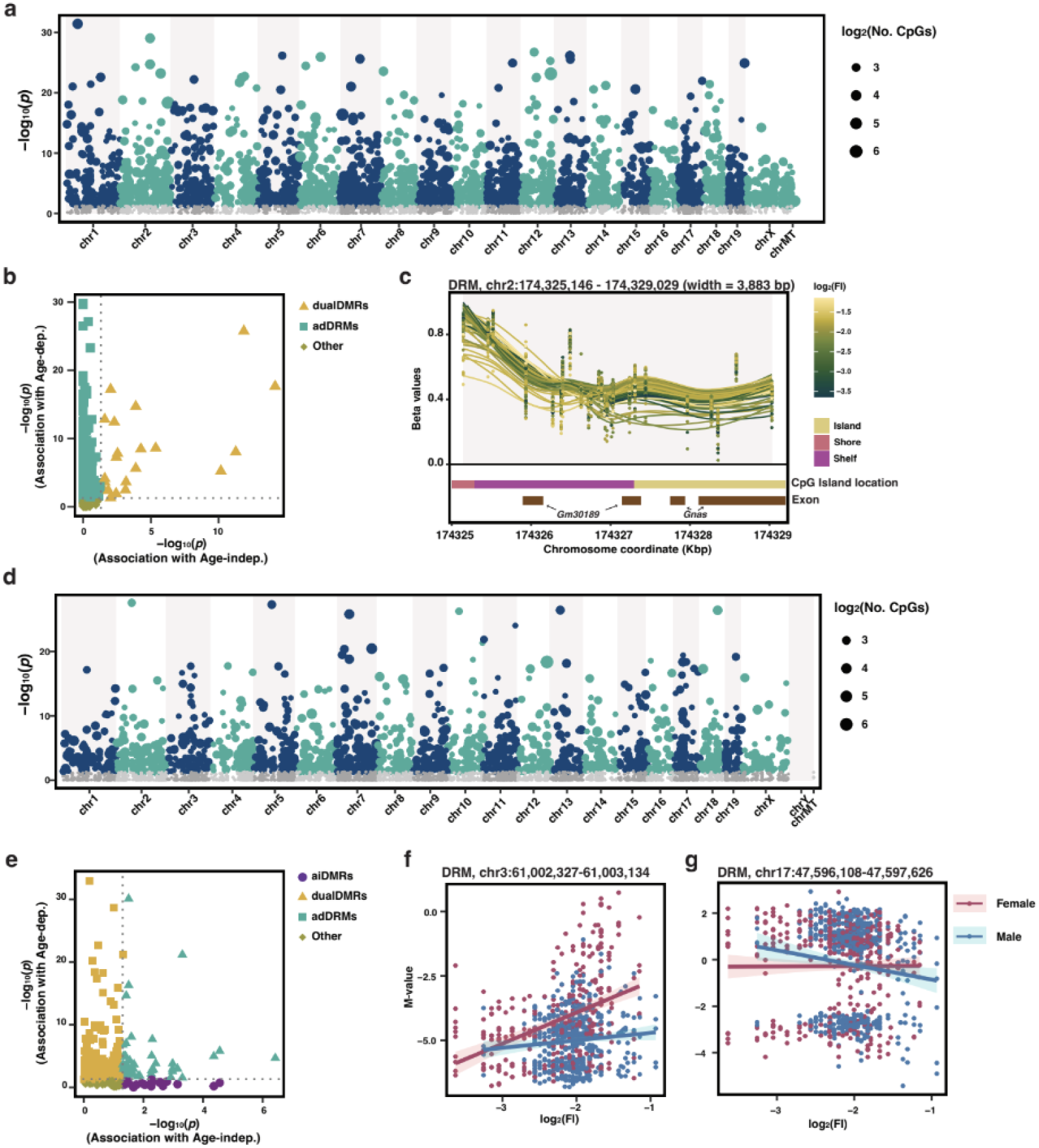
Visualization and characterization of DMRs in females and males. **a.** Manhattan plot showing the differentially methylated regions (DMRs) associated with frailty in females. DMRs were identified by Stouffer’s Z-score method and selected based on a *p*-value threshold of 0.05. The size of each point corresponds to the number of CpGs included. **b.** Scatter plot showing *p*-values of association between DMRs with age-indep. and age-dep. frailty in females. **c.** Illustrative examples of differentially methylated regions. Each point represents the methylation beta value of a CpG within the genomic location and CpGs from the same sample are connected by smoothed lines fitted using the LOESS method. Both points and lines are colored according to the level of log-transformed frailty index (log_2_FI). Distribution of CpG location categories (colored according to the category) and exons is presented in panels. **d.** Manhattan plot showing the differentially methylated regions (DMRs) associated with frailty in males. **e.** Scatter plot showing *p*-values of association between DMRs with age-indep. and age-dep. frailty in males. **f.** and **g.** Examples of DMRs that present sex-specific associations with frailty.

**Supplementary Fig.6.**
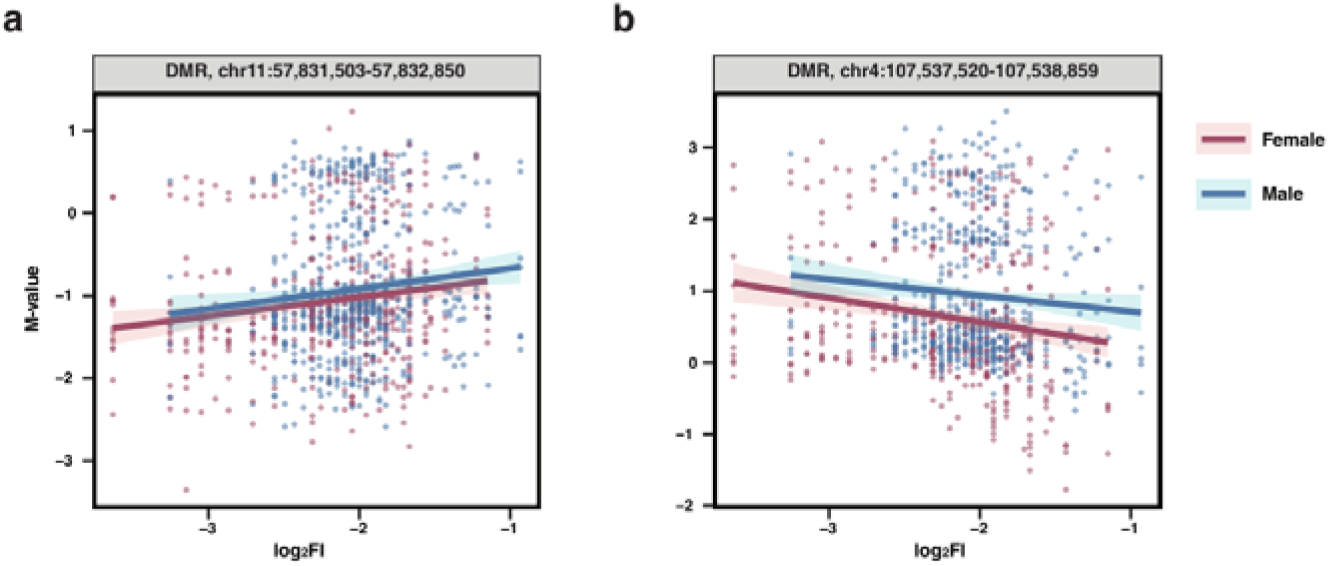
Examples of DMRs that present positive and negative effect sizes in association with frailty.

**Supplementary Fig.7.**
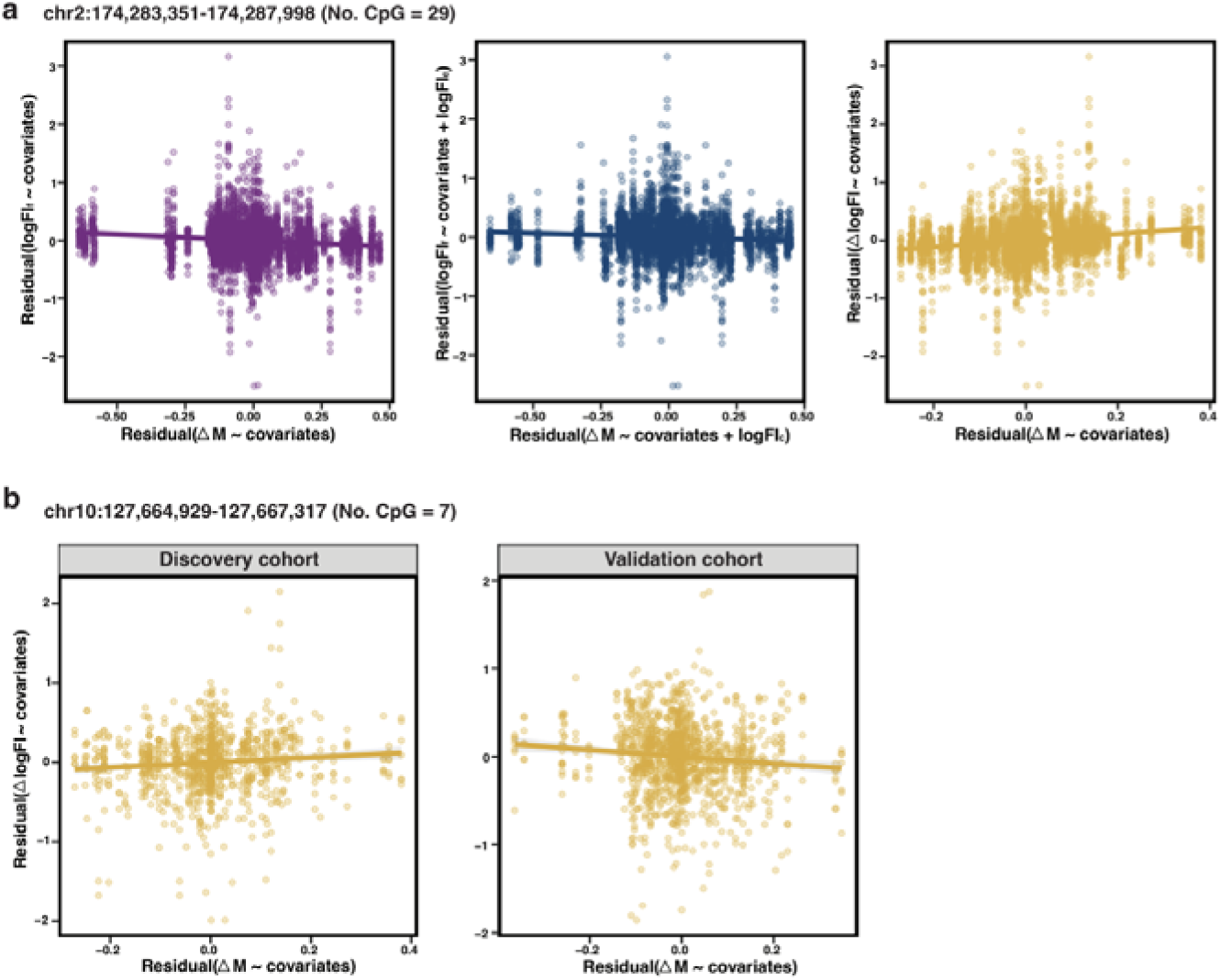
Partial regression plot showing example frailty-related DMRs association with frailty outcomes. **a.** The DMR presents opposite directions when associated with two future frailty outcomes. M-values of or changes of M-value from a prior time point (□M) for each CpG within the DMR were used as the independent variable and outcomes included logFI_c_ (log-transformed Frailty Index at Age_c_), logFI_f_ (log-transformed FI at a future age) and □logFI (the difference between logFI_f_ and logFI_c_). For logFI_f_, when used as the outcome to be associated with ΔM, further adjustment for logFI_c_ was performed. **b.** The DMR presents opposite directions when associated with □logFI (ΔM as the variate) across the discovery and validation cohorts.

**Supplementary Fig.8.**
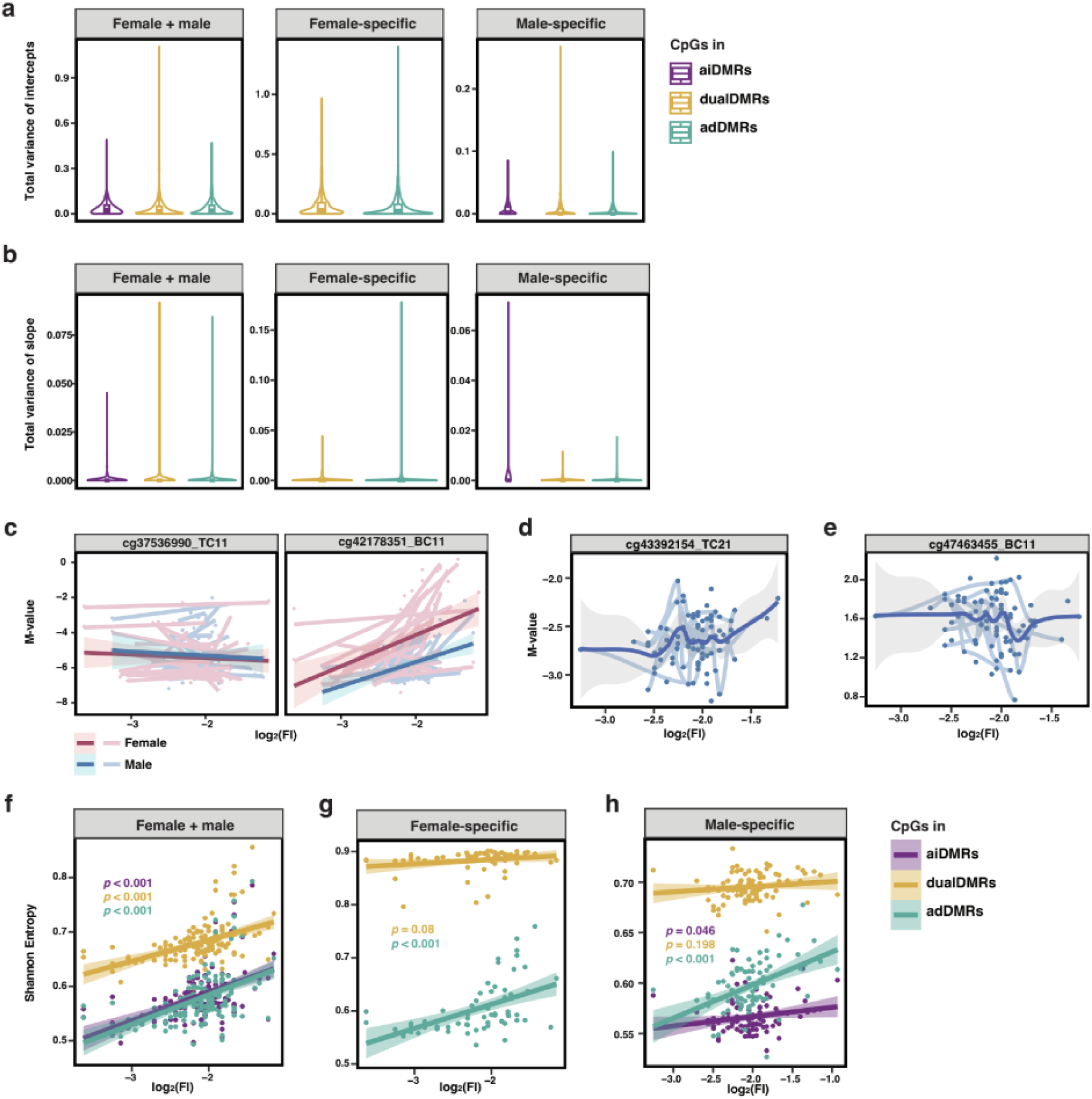
Variability analysis of frailty-related CpGs. **a.** Violin plot showing the total intercept variance stratified by the DMR categories the CpGs belong to in sex-inclusive and sex-specific groups. Variance was estimated using linear mixed models with random intercepts and slopes grouped by individual mice. **b.** Violin plot showing the total variance of slopes stratified by the DMR categories the CpGs belong to in sex-inclusive and sex-specific groups. **c.** Examples of DMRs that present similar slopes but distinct intercept in individuals. **d.** An example of DMRs that present deterministic methylation changes in frailty progression. **e.** An example of DMRs that present stochastic methylation changes in frailty progression. **f-h.** Scatter plots showing Shannon entropy of CpGs belonging to different DMR categories in frailty progression in sex-inclusive and sex-specific groups. Shannon entropy values were calculated using adjusted beta values, with sex effects regressed out, for each CpG within a DMR.

**Supplementary Fig.9.**
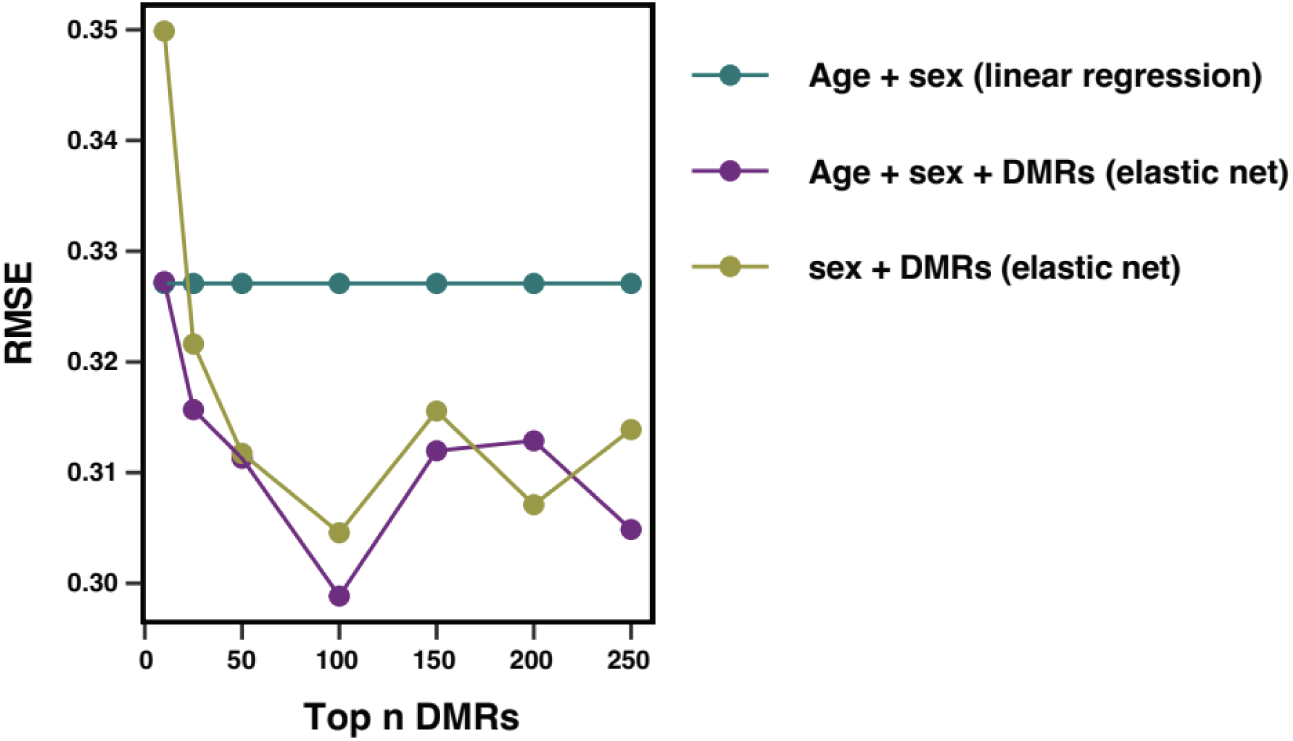
Comparisons of models that include diverse combinations of variables. The optimal number of top ranking DMRs (ranked by absolute effect size in association with frailty) were selected based on RMSE.

**Supplementary Fig.10.**
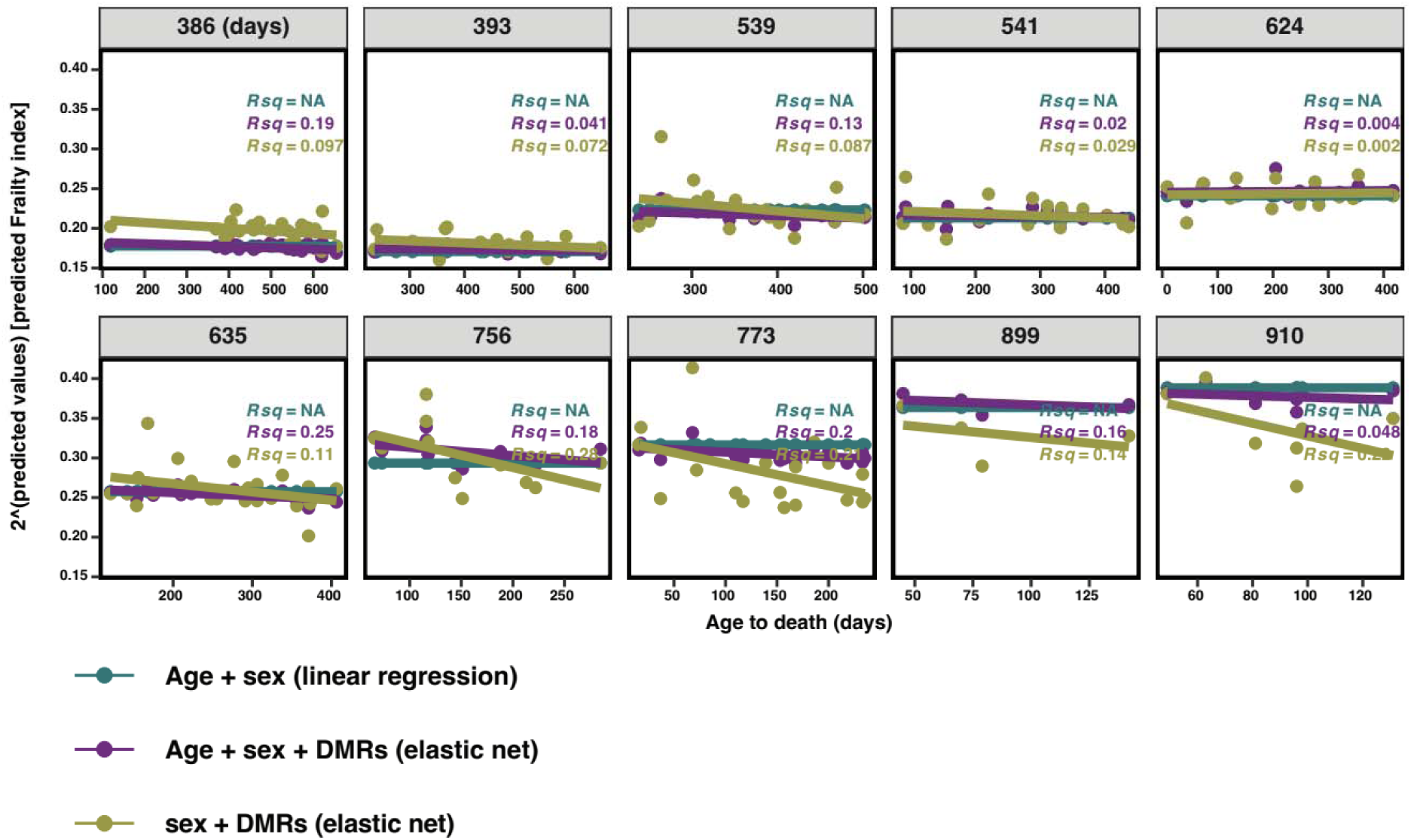
Comparisons of the performance of the epigenetic frailty clock and age+sex linear regression model in mice at the same age.

***Supplementary Tables1-9***

https://docs.google.com/spreadsheets/d/1gMrRaDpwGZ2D3VctHy9Cx-Pw1HuQMNE7_Sb1aOQD-Cg/edit?gid=109275884#gid=109275884

